# Identification of peptide coatings that enhance diffusive transport of nanoparticles through the tumor microenvironment

**DOI:** 10.1101/659524

**Authors:** Rashmi P. Mohanty, Xinquan Liu, Jae Y. Kim, Xiujuan Peng, Sahil Bhandari, Jasmim Leal, Dhivya Arasappan, Dennis Wylie, Tony Dong, Debadyuti Ghosh

## Abstract

In solid tumors, increasing drug penetration promotes their regression and improves the therapeutic index of compounds. However, the heterogeneous extracellular matrix (ECM) acts a steric and interaction barrier that hinders effective transport of therapeutics, including nanomedicines. Specifically, the interactions between the ECM and surface physicochemical properties of nanomedicines (e.g. charge, hydrophobicity) impedes their diffusion and penetration. To address the challenges using existing surface chemistries, we used peptide-presenting phage libraries as a high-throughput approach to screen and identify peptides as coatings with desired physicochemical properties that improve diffusive transport through the tumor microenvironment. Through iterative screening against the ECM and identification by next-generation DNA sequencing and analysis, we selected individual clones and measured their transport by diffusion assays. Here, we identified a net-neutral charge, hydrophilic peptide P4 that facilitates significantly higher diffusive transport of phage than negative control through *in vitro* tumor ECM. Through alanine mutagenesis, we confirmed that the hydrophilicity, charge, and their spatial ordering impact diffusive transport. P4 phage clone exhibited almost 200-fold improved uptake in *ex vivo* pancreatic tumor xenografts compared to the negative control. Nanoparticles coated with P4 exhibited ∼40-fold improvement in diffusivity in pancreatic tumor tissues, and P4-coated particles demonstrated less hindered diffusivity through the ECM compared to particles functionalized with gold standard poly(ethylene) glycol or iRGD peptide ligand. By leveraging the power of molecular diversity using phage display, we can greatly expand the chemical space of surface chemistries that can improve the transport of nanomedicines through the complex tumor microenvironment to ultimately improve their efficacy.

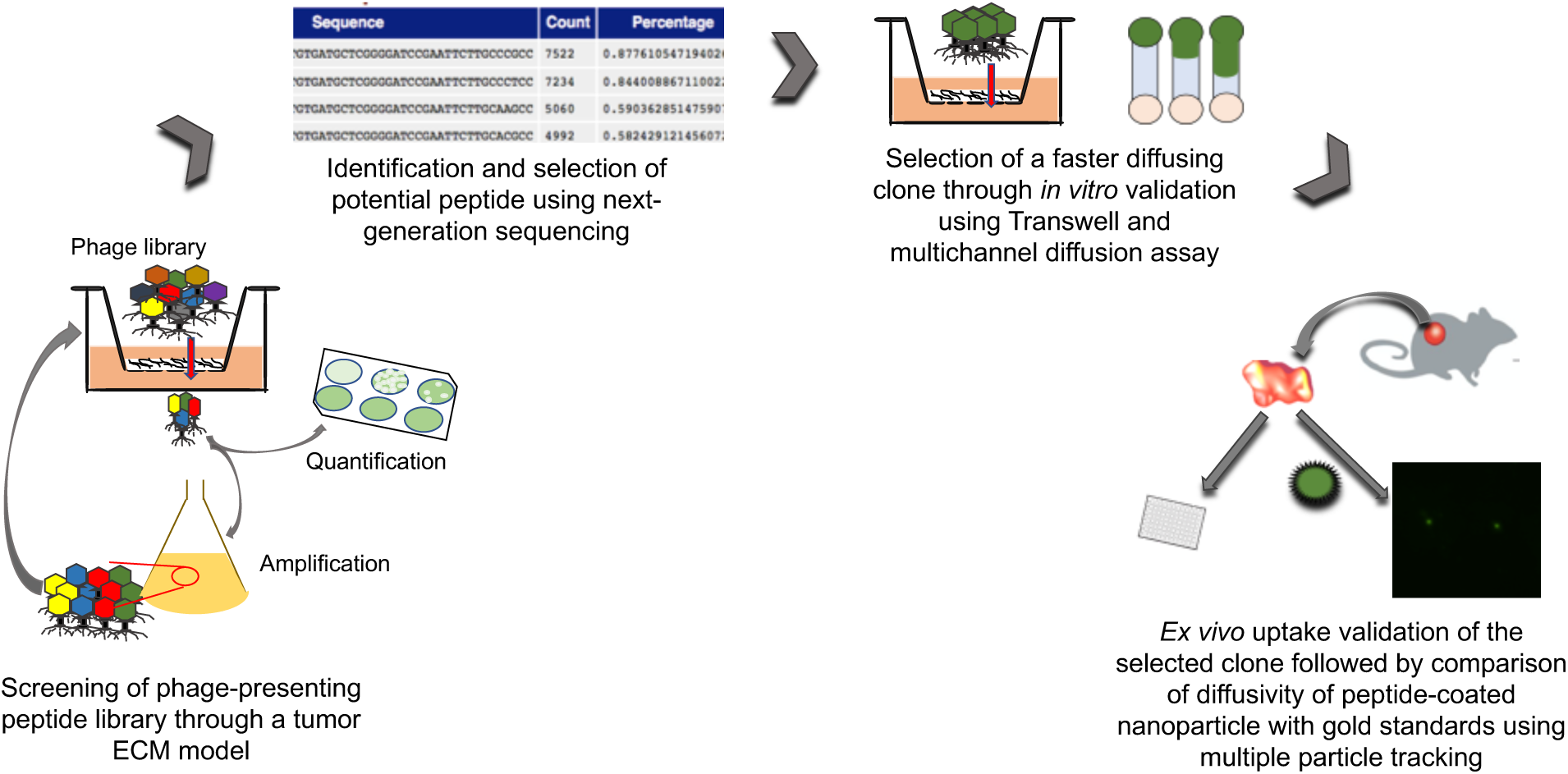

## Introduction

For successful delivery and efficacy of therapeutics in solid tumors, it is critical for drugs to reach as many target cells as possible to improve their therapeutic index and minimize drug resistance^1^. Increasing penetration of antibodies^2-4^, small molecule drugs^5–8^, and nanoparticles^9–12^ through the tumor microenvironment improves drug distribution, achieves tumor regression and significantly improves the therapeutic index of these compounds. The transport of these molecules through the heterogeneous tumor microenvironment significantly depends on their physicochemical properties and their interactions with the tumor extracellular matrix (ECM)^13^. The tumor ECM primarily composes of abnormal, overexpressed amounts of collagen, fibronectin, and glycosaminoglycans, which collectively form a net negative charged network that interacts with and impacts transport of molecules through size-based, hydrodynamic, and/or electrostatic interactions^14, 15^.

Nanoparticles are promising systems to deliver large quantities of therapeutic cargo while minimizing off-target toxicity in non-tumor tissues (reviewed in ^16^), and engineering these particles for effective transport through the tumor microenvironment is critical towards successful drug delivery. The physicochemical properties of nanoparticles, such as shape, size, and surface chemistry, govern their penetration into tumors, and optimization of these properties can enhance their transport for improved accumulation and retention. The tumor ECM acts as a steric barrier that can filter away particles based on size ^17, 18^. To address this problem, extensive efforts have been made to synthesize nanoparticles with smaller diameters ^10, 19–21,22–27^ or different non-spherical shapes ^19, 28, 29,30, 31,32^ to diffuse through the ECM mesh and improve penetration in select tumor types.

While the size and shape of nanoparticles have been engineered to circumvent the steric barrier of the ECM, molecules smaller than the pore size of the tumor ECM remain unable to traverse the barrier; the ECM also acts as an interaction barrier that prevents particle transport due to intermolecular interactions^33^. Here, there has been less work to engineer and optimize the surface chemistries of particles to interact (or not) with the ECM for improved tumor penetration. Studies indicated that positively charged liposomes^34^ and poly(lactic-co-glycolic) (PLGA) nanoparticles^35^ improved penetration in tumors, whereas in other studies, negatively charged liposomes^36^ and gold nanoparticles^37^ exhibited improved penetration in spheroids and tumors. Typically, the surfaces of most nanoparticles are functionalized with poly(ethylene) glycol (PEG) to improve the systemic circulation half-life, but the PEG coating additionally provides a net neutral, hydrophilic surface that facilitates penetration through the tumor ECM due to its inertness. PEG-coated PLGA and polystyrene nanoparticles exhibited enhanced diffusive transport and penetration through ECM gels^33, 38^ and tumor tissues^39–42^ than uncoated positive and/or negative charged particles. While PEGylation has been an attractive strategy for tumor penetration, there are some challenges that limit its feasibility. PEGylation has been shown to hinder cell internalization^43–45^ and endosomal escape^46^ needed for drug and gene delivery. Numerous studies have also shown that people possess pre-existing anti-PEG antibodies, which reduces the efficacy and clinical feasibility of PEG-conjugated therapeutics^47–50^. In addition, repeated systemic dosing of PEGylated formulations can result in the development of anti-PEG antibodies in living subjects^51–53^. These findings suggest while PEGylation can improve tumor penetration, there are challenges that motivate the need to find alternative surface chemistries.

Peptides are attractive biological materials for surface modification of nanomedicines. Their length and sequence can be controlled^54^, and they are chemically complex molecules that impart diverse functionality due to the amino acid sequence and their functional side chain groups^55^. In nanomedicine, peptides have been traditionally thought of and used as ligands that are conjugated to improve cell targeting and internalization of small molecules, macromolecules, and nanoparticles^56, 57^. For example, peptides such as iRGD have been discovered through phage display as ligands that bind tumor vasculature and target tumor cells^58, 59^. However, there has also been an increasing emphasis to exploit the chemistry of the amino acids and their physicochemical properties to develop peptides as surface coatings tuned for desired functionalities. For example, there has been extensive work to develop zwitterionic peptides with positive and negative charged amino acids as nonfouling coatings on surfaces^54, 60–62^. Additional work has showed that peptides with block configuration of charged amino acids (e.g. KKKKEEEE) can facilitate penetration through mucus hydrogels due to partitioning effects^63^.

In this work we use phage display technology as a combinatorial approach to generate and identify peptides as surface coatings to achieve improved diffusive transport through the tumor microenvironment. Phage, or bacterial viruses, can be genetically engineered to display peptides on their protein coat and can be pooled together to generate a collection of peptide-presenting phage libraries with excellent molecular diversity (i.e. 10^5^ – 10^9^ different peptide sequences)^64^. Specifically, we used T7 phage as the scaffold to display and identify peptide coatings. T7 is an icosahedral virus that can be engineered to display 415 copies of a peptide on its gp10 protein coat and has size features (∼70 nm diameter) on the scale of conventional synthetic nanoparticles. T7 phage are biological nanoparticles that can multivalently display different “peptide-based” surface chemistries that can be combinatorially screened to identify peptide coatings. Here, we engineered random peptides to be displayed on T7 phage and screened the library against the selective pressure of ECM components found in solid tumors to collect peptide-presenting phage that permeated through the ECM. After successive rounds of selection, permeating phage were harvested, and using next generation sequencing and analysis, the most frequent peptides and motif sequences were identified and subsequently validated in transport studies. Diffusion assays were done to measure and identify peptide-presenting phage that exhibited most unhindered diffusivity through the ECM. Using one of the selected peptides, P4, as a model, we performed site-directed alanine mutagenesis to study the role of hydrophilicity, charge, and the order of the amino acid sequence on diffusive transport through the ECM. We next measured the ability of select peptide-presenting phage to achieve uptake in *ex vivo* ECM-rich pancreatic cancer xenografts. To demonstrate its feasibility as a surface coating for synthetic nanoparticles, the tumor penetrating peptide was conjugated onto polystyrene nanoparticles, and their diffusive transport in pancreatic tumor tissue was measured using multiple particle tracking and compared to PEG-functionalized and control uncoated nanoparticles. Phage display provides the molecular diversity that can be leveraged to screen and select peptides as surface coatings that facilitate diffusive transport and establish that charge, hydrophilicity, and their orientation govern transport through the tumor ECM. This work highlights the promise of peptides as alternative surface chemistries to improve transport and delivery of nanomedicines for solid tumors and potentially other disease microenvironments.

## Results and Discussion

The tumor ECM is a barrier that restricts diffusive transport of therapeutics. The interaction between the ECM with the surface physicochemical properties of nanomedicines dictate their transport through this barrier. Here, the objective was to identify peptides as surface coatings using combinatorial selection against the tumor ECM and validate that they improve diffusive transport through the ECM. A second objective was to understand how physicochemical properties of peptides, including charge and hydrophobicity, can affect transport through the ECM barrier.

### Screening of peptide-presenting phage libraries against tumor ECM model

We used combinatorial screening (**Figure 1a**) with next-generation sequencing and analysis to identify peptides that can achieve diffusive transport through the ECM network present in tumors. A cysteine constrained 9-mer random peptide-presenting T7 phage library was engineered to effectively serve as a collection of peptide-based surface chemistries with a combination of amino acids possessing varying physicochemical properties. The phage library was incubated with a transwell coated with ECM components collagen I (7.8 mg/ml) and hyaluronic acid (HA) (0.84 mg/ml), which are predominant in the tumor microenvironment of pancreatic cancers ^65–67^. The charged ECM components serve as an electrostatic interaction filter that has been shown to hinder diffusion of charged molecules ^15, 67^. A 1 mm thickness of tumor ECM was prepared in the insert of a 24-well transwell. The phage library was added at the top of the ECM to allow it to diffuse through the tumor ECM model at 37 °C. After 1 hour of incubation, phage that permeated through the ECM was collected, and a fraction of the output was quantified by standard double-layer plaque assay. The remaining fraction was amplified in *E. coli* to make more copies needed for the subsequent round of screening. This screening was iterated for three additional rounds and the overall selection was performed in replicate. Selection of peptides using phage libraries is typically reported from one or two screenings^68, 69^.

**Figure 1.**
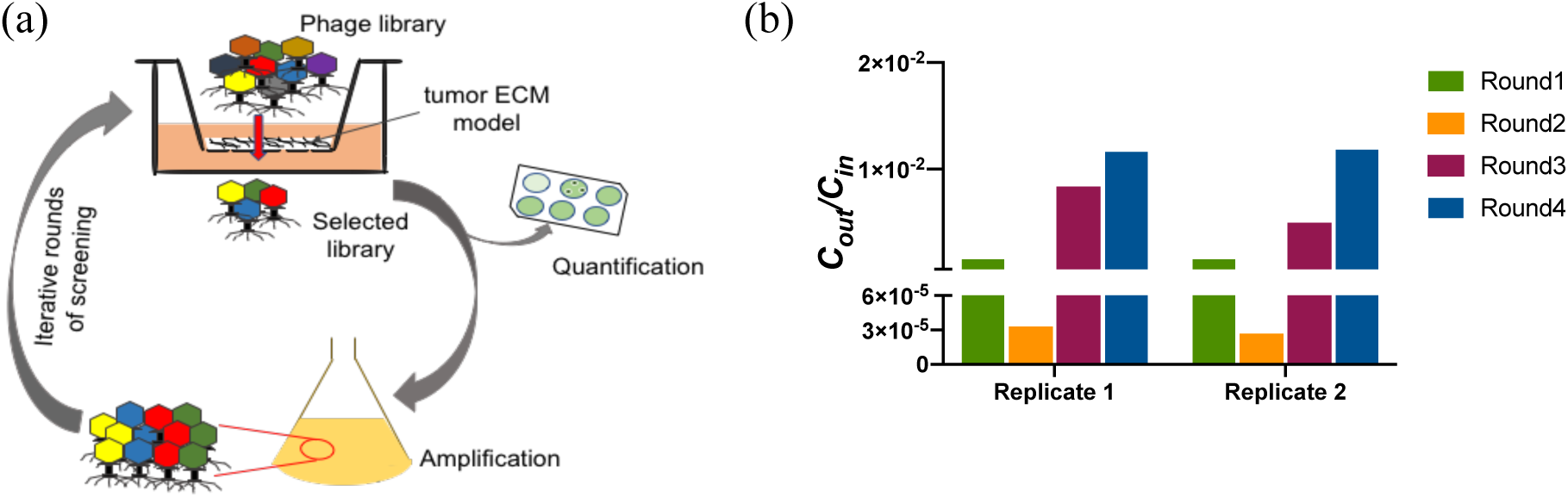
**(a)** Schematic of peptide-presenting phage library screening through a tumor ECM model. Phage library was allowed to diffuse through the tumor ECM model prepared in the insert of a transwell. After one hour, phage was collected from the receiving chamber and part of the eluate was used for quantification using double-layer plaque assay. The remaining fraction was amplified for use in subsequent screening, which was performed for a total of four rounds. **(b)** Enrichment of phage library diffusing through in vitro tumor ECM. *C*_out_ represents the output titer of the phage library after 1 hour of permeation through the tumor ECM, and *C*_in_ represents the input titer. Round1, Round2, Round3 and Round4 signify the iterative rounds of phage library screening. Replicate 1 and Replicate 2 are the two replicates of the screening.

Iterative screening allows the tumor-like ECM to serve as a selection pressure to filter away phage unable to consistently penetrate through the ECM, irrespective of the heterogeneity of its mesh network. In each replicate, we observed an increased amount of phage that permeated through the tumor-like ECM after four rounds (**Figure 1b**). The average library titer (i.e. amount of phage) of both replicates increased 200-fold and 400-fold in the third and fourth round compared to the second round. In the third and fourth rounds, the increase of permeating phage suggests there is enrichment of the library for phage that may possess desired functionalities (i.e. unhindered transport) through the tumor-like ECM. There was no increase in the first two rounds, but the lack of initial enrichment is typically observed in phage selections^70^. The enrichment of phage during selection is expected, as observed with affinity-based panning against specific targets ^70–72^.

### Identification and selection of peptides using next generation sequencing (NGS)

From each round of panning, a fraction of the permeated phage was subsequently used to identify the peptide sequences. Since the DNA sequence of the individual phage encodes for the peptide displayed on the phage surface, we used NGS to obtain a high number of DNA reads not feasible with Sanger sequencing, and the sequences were translated and sorted to identify abundant peptides present during our selection. DNA from the eluted phage was isolated and PCR amplified for the region encoding the random peptides, and the DNA were individually barcoded to identify selection round and replicate. The pooled, barcoded DNA sequences were run on Illumina MiSeq to obtain up to 10^5^−10^6^ reads per sample. Using customized Python and bash scripts, the DNA reads were trimmed to the DNA sequences that encode for the displayed peptide and translated, and the peptides were counted and sorted by their frequency relative to the total number of reads. Sequences that contained truncations, premature stop codons, and/or did not contain the full amino acid sequence were excluded in subsequent analysis and experiments. We then determined the overall physicochemical properties of the thirty most frequent peptide sequences by *in silico* analysis (**Figures S1a and S1b**).

From the thirty most frequent peptides after four rounds of selection, we selected five peptide-presenting phage to subsequently be validated in transport studies. Three peptides known as P1, P2 and P3-peptides were selected based on their frequency and reproducibility, i.e. if they are present within the thirty top-frequent sequences of both replicates. The frequency of each individual peptide as a percentage of total number of reads from each round of panning was quantified (**Figure 2**) to determine if phage-displayed peptides were enriched during selection against the tumor-like ECM. In general, the number of the selected peptides were enriched over the four rounds of screening. The relative abundance of the individual peptides did not increase from original library (R0) to first round of screening (R1), but they are enriched 10 to 40-fold from round 1 (R1) to round 2 (R2) and had moderate increase in rounds 3 and 4 (R3 and R4); these results have been observed by others and is indicative of their enrichment^69–71, 73^. From NGS data, we selected one peptide sequence from the original, naïve library as a negative control for our subsequent experiments. This control peptide sequence is present in the original library, but its frequency decreased with successive rounds of selection, which suggests it was unable to withstand the selection pressure of the tumor-like ECM and cannot permeate through the barrier.

**Figure 2.**
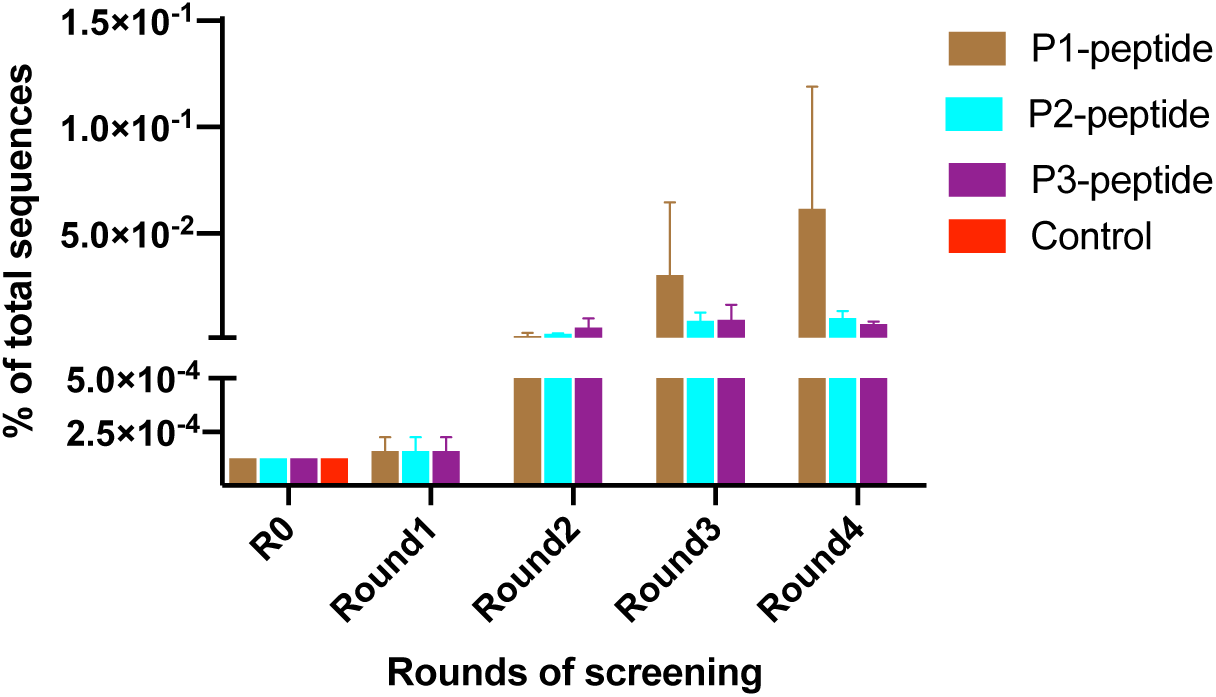
Sequences selected based on their frequency and reproducibility (P1, P2, and P3) were enriched over the rounds of screening. The negative control selected from the naïve library did not withstand the selection pressure and decreased in relative frequency with successive rounds of selection. R0: Original naïve library, P: Peptides selected from phage display. Data represents mean ± SD of two replicates of library screening.

Two additional sequences, P4 and P5-peptides, were selected from multiple sequence alignment using Seq2Logo as described elsewhere^69, 74^. Here, multiple sequence alignment of the top thirty frequent peptides from the fourth round of both replicates was done to identify the most abundant amino acid at each position of the 7-mer^75^. From this analysis, we generated and selected synthetic motifs P4-peptide from replicate 1 and P5-peptide from replicate 2 for subsequent studies.

The selected peptides along with their physicochemical properties (net charge at pH 7.0 and hydropathy/hydrophobicity) are listed in **Table 1**. Three of the five sequences selected are hydrophilic, one is hydrophobic, and one of the sequences is neither hydrophilic nor hydrophobic, which we term as hydro-neutral. The net charge of two of the selected sequences is negative, two are positive, and one of them has a net neutral surface charge at pH 7.0. The net charge of all the selected peptides are near neutral, ranging from −1.0 to +1.1.

**Table 1.** Peptides selected for validation based on frequency, reproducibility and multiple sequence alignment of the thirty most frequent sequences from both replicates of phage library screening, along with their physicochemical properties.

| Peptide Name | Hydropathy | Net charge at pH 7 |
| --- | --- | --- |
| P1-peptide | -1.2 | +1.1 |
| P2-peptide | +1.0 | -1.0 |
| P3-peptide | 0 | -1.0 |
| P4-peptide | -0.7 | 0 |
| P5-peptide | -0.1 | +1.0 |
| Control-peptide | -0.7 | -5.0 |

The selected and control sequences were each genetically engineered into the T7 vector to display the peptides on the surface of the T7 phage (denoted as P1 to P5-phage, insertless (WT), and control-phage). The T7 phage are biological nanoparticles; the peptide-presenting phage clones have nanoscale dimensions, and in addition, phage have served as a template for molecular assembly of nanoscale materials and have been used in various applications in nanomedicine^76^. The diffusion coefficient of clones is representative of the unhindered diffusivity of individual clones in phosphate buffered saline (PBS) using dynamic light scattering (**Table S1**). The measured diffusion coefficient of the insertless, WT phage (hydrodynamic diameter ∼70 nm) correlates with its theoretical calculation following the Stokes-Einstein equation.

### Improved diffusion of selected peptide-presenting phage through *in vitro* ECM

Next, we confirmed that selected and motif peptide-presenting phage exhibited improved diffusion through the tumor-mimetic ECM compared to control phage. Here, we measured the diffusion of the peptide-presenting phage through tumor-like ECM using a fluorescence-based microfluidic channel diffusion assay, as described elsewhere ^66^. We prepared the ECM within the channel of a microfluidic chamber µ-slide (**Figure 3a**). The tumor ECM in the channel provides a semi-infinite barrier to test the diffusivity of the phage clones. The encapsulated DNA of the phage was fluorescently labeled to avoid surface labeling of phage, which could change the surface properties of the phage and affect interaction and transport through the ECM. Phage were added to the input reservoir, and an equal volume of PBS is added to the output reservoir to maintain the hydrostatic pressure and to achieve a concentration gradient through the channels. Fluorescent images of the channel were acquired at different time points and processed to identify the fluorescence intensity at each time point over the distance of the entire channel (**Figure 3b**). The normalized intensity value over the channel length and the entire diffusion time was used for non-linear curve fitting with the Fick’s second law of diffusion to calculate the diffusion coefficient of each clone using a customized MATLAB script (**Figure S2**).

**Figure 3.**
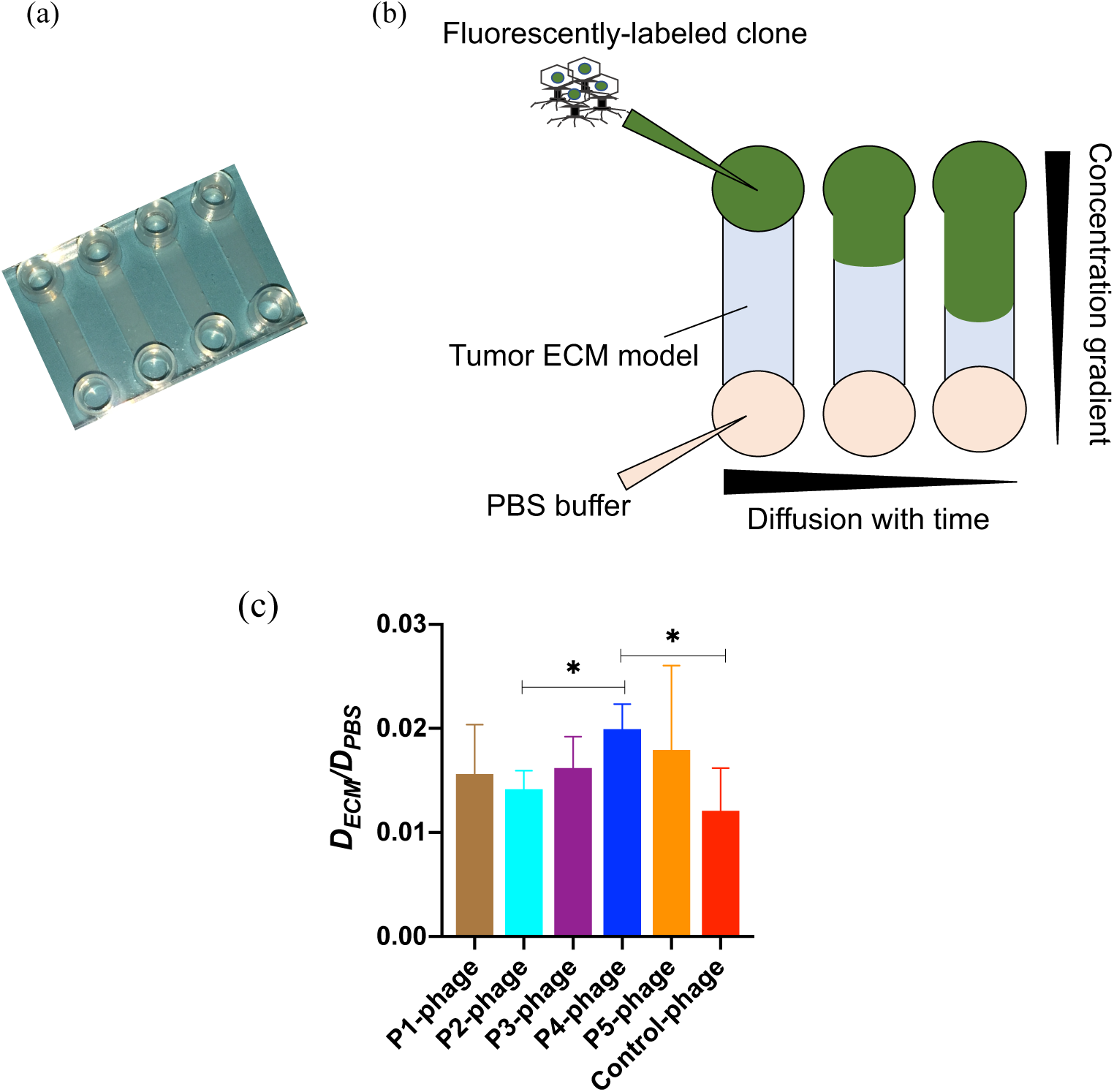
(a) Tumor ECM-loaded µ-slide (b) Schematic representation of the diffusion of fluorescently-labeled phage clones through the tumor ECM along the length of the channel and with time (c) The diffusion coefficient through ECM normalized to the theoretical unhindered diffusivity in PBS of selected clones is compared to the negative control-phage in the microfluidic channel. Data represents mean ± SD (n=3), unpaired, two-tailed Welch’s t-test (p<0.05).

The diffusivity of the clones through the ECM varied from 6.40×10^-14^±2.17×10^-14^ m^2^/sec for the negative control-phage to 1.05×10^-13^±1.27×10^-14^ m^2^/sec for P4-phage, which was in the same order of magnitude as the diffusivity of similar sized T4 phage through gels and artificial biofilm models^77, 78^. Comparable values of 1.62×10^-12^ m^2^/sec and 3.39×10^-13^ m^2^/sec were reported for diffusion of 50 and 120 nm gold nanoparticles through a similar microchannel with collagen I^66^. To determine how the ECM affects diffusion, we calculated the effective diffusivity of the clones through the tumor-like ECM with respect to the theoretical diffusivity of the WT in PBS (**Figure 3c**).

Diffusion of the negative control was hindered 80-fold through the dense tumor ECM. However, all the selected clones exhibited improved diffusivity through the tumor-like ECM compared to the control (**Figure 3c)**. Since the only difference between the selected clones and the control was the sequence of the displayed peptides, this finding suggests that the physicochemical properties of the displayed peptide likely impacted binding to the ECM and subsequent diffusion. In particular, the P4-phage demonstrated significantly higher diffusivity through the ECM compared to the control-phage. The hydrophilic and net neutral charged peptide sequence P4 was selected for subsequent studies since it demonstrated highest average diffusivity through the *in vitro* tumor ECM in the microfluidic and transwell diffusion studies (**Figure 3c** and **Figure S3**).

### Effect of specific physicochemical properties on the diffusivity is investigated by site-directed alanine mutagenesis

While net-neutral charge, hydrophilic polymer coatings have improved transport of nanocarriers through ECM ^33, 39, 41^, the polymer coating consists of a single repeating unit (e.g. H-[O-CH_2_-CH_2_]_n_-OH for PEG). Alternatively, due to molecular diversity of amino acids and their ordering in the peptide sequence, we can systematically manipulate select amino acids and determine their contribution to peptide-mediated transport through the ECM. To determine the impact of charge, hydrophobicity, and spatial order of physicochemical properties on transport through the ECM, we performed alanine mutagenesis on the selected P4. We substituted specific amino acids with alanine, which has an inert methyl functional side chain group that does not change the main conformation of the peptide, and thereby we removed the functional groups to create mutants with different surface physicochemical characteristics. We investigated the effect of net charge, hydrophobicity, and order of the amino acid sequence on diffusive transport through the tumor ECM using the aforementioned microchannel diffusion assay.

The P4 sequence has one basic amino acid, and the remaining residues are polar amino acids. To alter the net charge, we mutated the basic and acidic positions to alanine to create negative (−1) and positive (+1) charge modifications, respectively (**Table 2a**). The diffusivity of the original and charge modified clones through tumor-like ECM was measured, and their diffusivity was compared to the theoretical diffusivity of WT in PBS. The original neutral clone exhibits approximately 1.5-fold higher effective diffusivity compared to the positive and negative modifications (**Figure 4a**). As a result of the alanine mutations, the hydrophobicity of the sequences has also been changed due to the removal of the polar acidic and basic amino acid residues. A similar decrease in hydrophilicity was also observed during library screening, where highly positive and negative charged peptides (with polar basic and acidic residues) were filtered out with increased rounds (**Figures S1a** and **S1b**).

**Table 2a.** Alanine mutagenesis to study the effect of net surface charge.

| 1. Effect of Surface Charge |  |  |  |  |  |  |  |  |  |
| --- | --- | --- | --- | --- | --- | --- | --- | --- | --- |
| Sequences | Aminoacid positions in peptide sequence |  |  |  |  |  |  | Hydropathy | Net charge (pH 7) |
|  | 1 | 2 | 3 | 4 | 5 | 6 | 7 |  |  |
| P4-Neutral |  |  |  |  |  |  |  | -0.756 | 0 |
| Negative | A |  |  |  |  |  |  | -0.122 | -1 |
| Positive |  |  |  | A |  |  |  | -0.167 | 1 |

**Table 2b.**
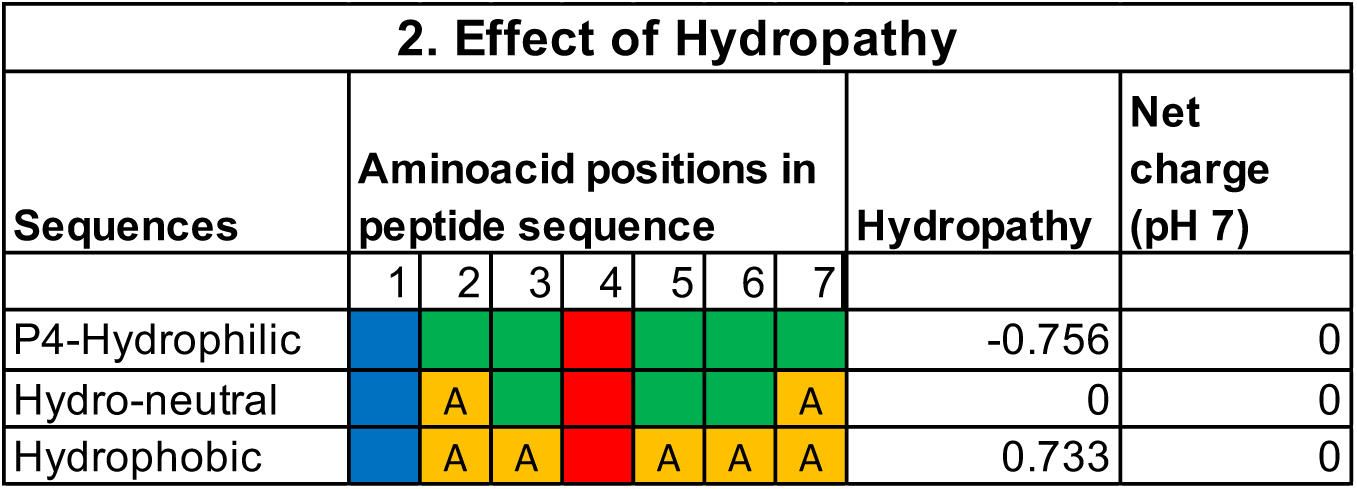
Alanine mutagenesis to study the effect of hydropathy.

**Table 2c.**
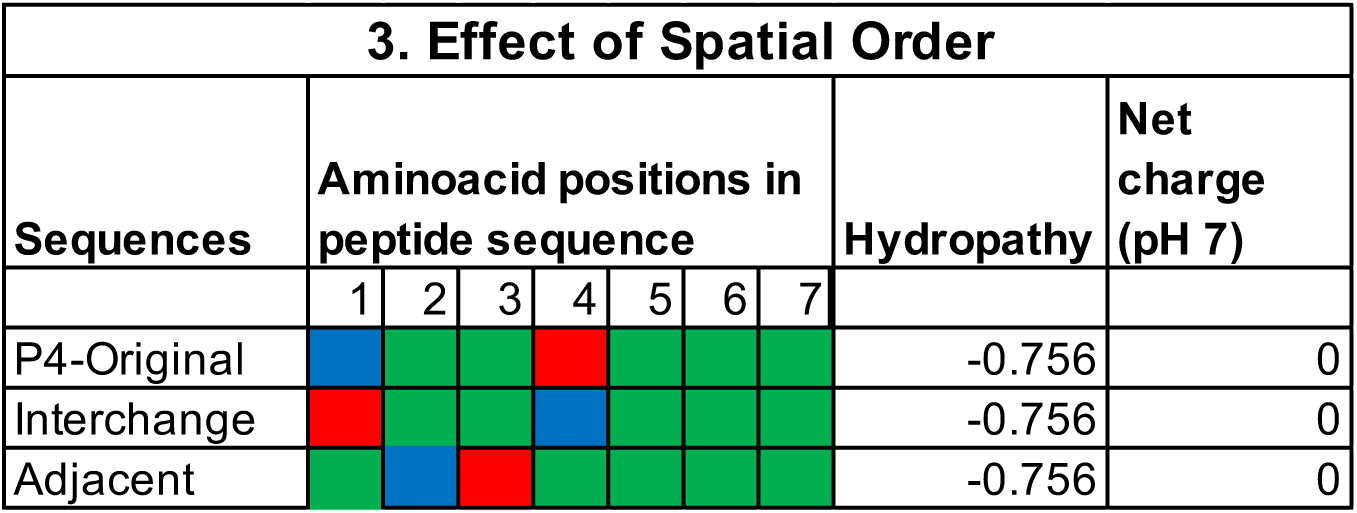
Alanine mutagenesis to study the effect of spatial order of the amino acids.

**Figure 4.**
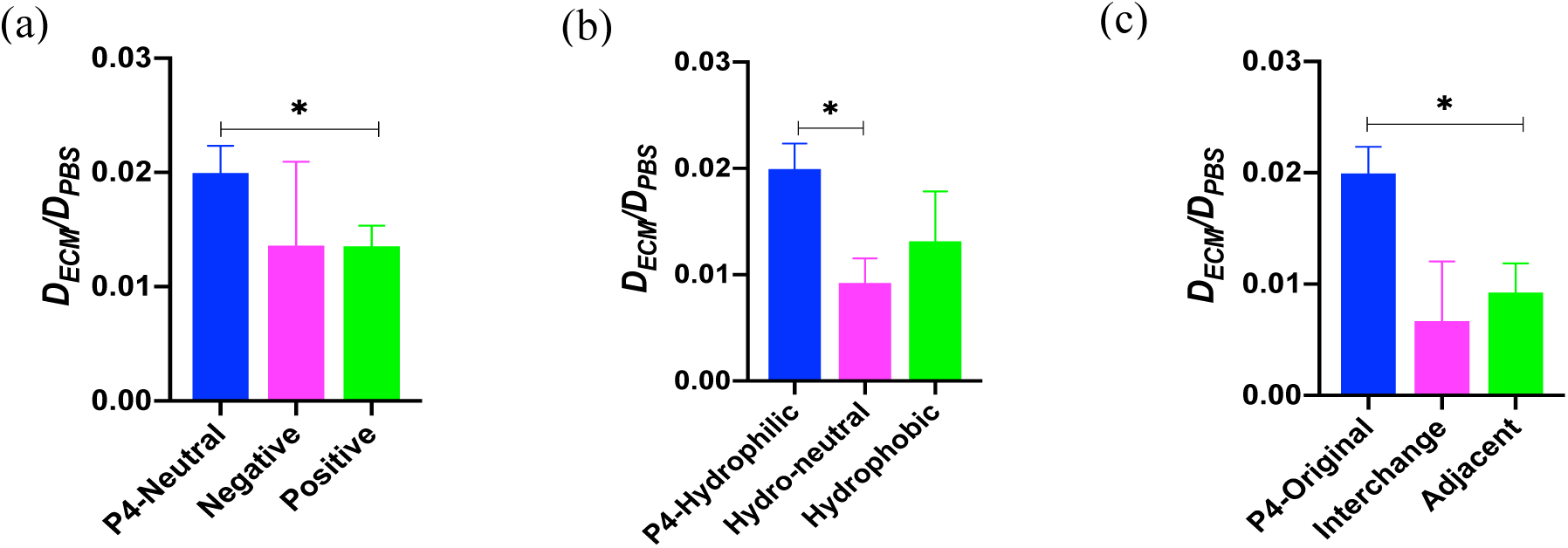
Alanine mutagenesis. Native and mutant P4-phage were used in microfluidic diffusion assays to understand the effect of (a) surface charge, (b) hydrophobicity, and (c) spatial order of the amino acids on diffusion through ECM (D_ECM_) relative to their unhindered diffusion through PBS (D_PBS_). Hydro-neutral signifies neither hydrophilic nor hydrophobic. Data represents mean ± SD (n=3), One-way analysis of variance (ANOVA) with Dunnett’s T3 multiple comparison (p<0.05).

Next, we determined the role of hydrophobicity on transport of our selected peptide through the ECM. We modified the hydrophobicity of the original sequence through alanine mutagenesis of some or all of the polar amino acids, as indicated in **Table 2b**. In one of the modifications, we designed a peptide that is neither hydrophilic nor hydrophobic, which we termed hydro-neutral. We also generated a hydrophobic sequence where all of the polar amino acids were substituted with alanine. These modifications retained the same net charge of the original sequence while altering the hydrophobicity (**Table 2b**). The original hydrophilic clone demonstrated 2-fold and 1.5-fold higher diffusivity than the hydro-neutral and hydrophobic clones, respectively (**Figure 4b**). This result is similar to reports where hydrophilic PEG coatings on hydrophobic polystyrene and poly(lactic-co-glycolic) nanoparticles improved their diffusivity through brain^41^ and breast^39^ tumor microenvironments. Hydrophobic surfaces of particles can strongly bind to the ECM^41^, and hydrophilic coatings minimize adhesion.

Next, we made alterations to the order of the amino acid sequence to understand how changing the spatial order of the physicochemical properties affects diffusive transport through the tumor ECM. We made two modifications—(1) interchange the position of the basic and the acidic amino acid residue of the P4 sequence and (2) place the acidic and basic residues adjacent to each other (**Table 2c**). The net charge and hydrophobicity of the mutant sequences remain the same as the original sequence. From diffusion assays, the original selected sequence exhibits three-fold and two-fold higher effective diffusivity compared to the interchanged and adjacent mutants, respectively (**Figure 4c**). This result suggests that the order of the amino acid sequence significantly impacts the diffusivity through the tumor-like ECM. Others have previously shown that modifying the spatial configuration of charge modulates the transport of peptides through a mucin hydrogel^63^. There, two peptides with the same net charge but different amino acid configuration had different transport behavior through mucin; a peptide possessing block charge (e.g. KKKKEEEE) demonstrated partitioning and a sharper diffusion gradient than alternating amino acids with same composition and net charge (e.g. KEKEKEKE)^63^. Placing the basic and acidic amino acids adjacent to each other neutralizes the effect of the charged amino acids, thereby altering the interaction with the surrounding environment^63^. However, it is unknown why there is a decrease in the diffusivity through the tumor ECM by interchanging the position of the charged amino acids, and further studies are needed to understand their interaction with the ECM and correlate to transport.

### Higher equilibrium uptake of selected peptide in *ex vivo* pancreatic tumor tissues

Building on our *in vitro* studies, we next validated that our selected clone P4-phage improved transport and uptake in tumors compared to the negative control-phage. We measured and quantified *ex vivo* uptake of our peptide-presenting phage in ECM-rich BxPC3 pancreatic tumor xenografts^79^ as described by Bajpayee et al.^80^. Briefly, the BxPC3 tumor tissue slices were preequilibrated in PBS with protease inhibitor and incubated with in a bath of sample, P4 or control-phage. We calculated the clone uptake ratio in the tissue by quantifying the amount of phage in the tissue to the amount in the incubation bath over several days. The average uptake of selected clone P4-phage dramatically increased from 1.67×10^3^±2.08×10^3^ after 1 day to 1.34×10^8^±2.31×10^8^ after 5 days of the incubation, whereas the tumor uptake of the negative control-phage only increased from 1.12×10^3^±4.94×10^2^ in 1 day to 7.27×10^5^±1.32×10^5^ in 5 days (**Figure 5**). Consequently, the uptake ratio of P4-phage is 200 times higher than that of the negative control-phage after five days of incubation with the *ex vivo* tumor tissue. At initial timepoints, the rate of uptake is similar for both clones, but at later timepoints, there is a larger concentration gradient for our selected clone. This gradient allows continued diffusion and further transport of the P4-phage into the tumor tissue than observed with the control-phage. The difference in concentration gradients resulting in improved tissue uptake has been observed in cartilage explants ^80–82^ and mucus hydrogels ^63^.

**Figure 5.**
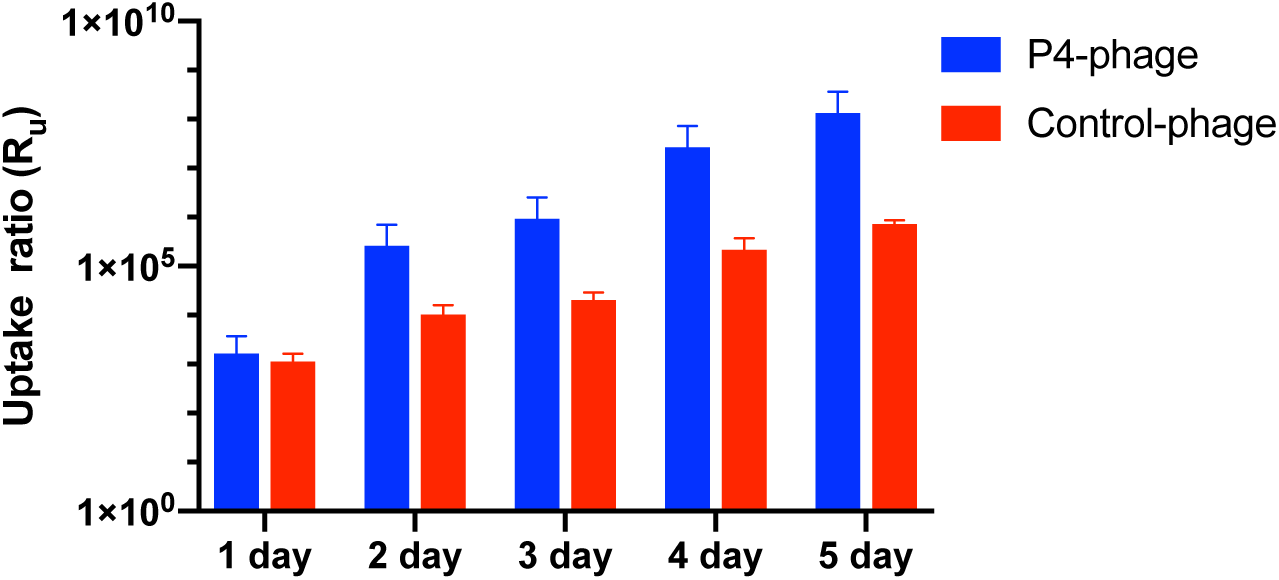
Uptake ratio of clones in *ex vivo* tumor tissue. Uptake ratio of the selected clone P4-phage compared to the negative control-phage in *ex vivo* tumor tissue over five days of incubation. Data represents mean ± SD (n=3).

### Improved transport of nanoparticles in tumor tissue with peptide coatings

While we have demonstrated that P4 peptide facilitates and improves transport of phage (P4-phage) through the tumors, we wanted to translate peptides from the structural context of phage and demonstrate their ability as surface coatings to improve diffusive transport of synthetic nanoparticles through the tumor microenvironment towards improving drug delivery. Here, fluorescent nanoparticles were conjugated with our selected peptide or gold standard controls from literature, and their displacement in tumors was tracked and measured using particle tracking microscopy.

We compared the penetration of P4-coated nanoparticle through the tumor ECM with two functionalized controls that have been previously used to improve the penetration and/or uptake of nanoparticles. iRGD peptide (CRGDKGPDC), which is a cysteine constrained 9-mer peptide sequence identified by phage display, has been extensively demonstrated to improve solid tumor penetration^59^. In addition, low molecular weight PEG surface modification of nanoparticles has been shown to improve their diffusivity through various biological barriers including mucus^83, 84^, brain extracellular space^41^ and solid tumor ECM^39^. We conjugated the peptides and PEG coatings to fluorescent carboxylate nanoparticles via standard EDC bioconjugation chemistry. We selected PEG 1 kDa to match with the size of the peptide used in this work. As a result, the size and surface properties of all the particles are similar (**Table 3**), and therefore any change in the diffusivity of the particles is due to their interactions with the tumor tissue.

**Table 3.** Characterization of the PEG/peptide coated nanoparticle formulations.

| Sample | Size (nm) | PDI | Zeta (mV) | <MSD><sub>PBS</sub>/<br><MSD><sub>tumor</sub><br>(1 sec time lag) |
| --- | --- | --- | --- | --- |
| Uncoated | 145.9±0.9 | 0.09 | -71.4±2.6 | 2373.63 |
| P4-coated | 152.5±2.9 | 0.08 | -59.9±3.9 | 57.24 |
| iRGD-coated | 153.1±3.0 | 0.08 | -62.9±3.3 | 137.66 |
| PEGylated | 150.1±0.2 | 0.09 | -52.3±3.4 | 238.64 |

We performed multiple particle tracking to measure the displacement of the surface modified nanoparticles through *ex vivo* BxPC3 tumor tissue xenografts (**Script S1**). The collective movement of uncoated nanoparticles was strongly hindered in the tumor tissue (**Figure 6** and **Table 3**). The average MSD of the uncoated particles was 10^-4^ to 10^-3^ µm^2^, which is nearly ten-fold less compared to the value reported by Dancy et al. using 100 nm beads (measured size of 96 nm) through *ex vivo* breast tumor slices^39^. The discrepancy can be attributed to the difference in the actual hydrodynamic size of the nanoparticles used in both studies and/or the heterogeneous composition and pore size distribution between the breast tumor versus pancreatic tumor microenvironments. The PEG and peptide coatings improved diffusion of the nanoparticles, as indicated by the upward shift in the ensemble-averaged mean squared displacement (MSD) at a time scale of 1 sec (**Figure 6a**). Both PEGylation and iRGD-coating improved the MSD of the nanoparticle 10- and 15-fold, respectively (**Figure 6b**). This improvement is most likely due to their hydrophilic and less negative surface charge properties. Interestingly, the P4-peptide coating improved the MSD of the nanoparticle 40-fold, exhibiting greater improvement than the other coatings. Examining the individual particle diffusivity binned together as ensemble measurements through the tumor ECM, there is a shift in particle distribution towards higher MSD upon PEG or P4-peptide coating (**Figures 6c and 6d**). In particular, there is a larger fraction of peptide-coated nanoparticles that are mobile and demonstrate rapid diffusivity (log MSD is greater than −0.5) compared to PEG-coated nanoparticles (5% to 1%, respectively). Given that the same molar ratio was used to conjugate peptide/PEG onto nanoparticles, this result suggests that the optimal peptide coating improves unhindered particle transport through the tumor ECM.

**Figure 6.**
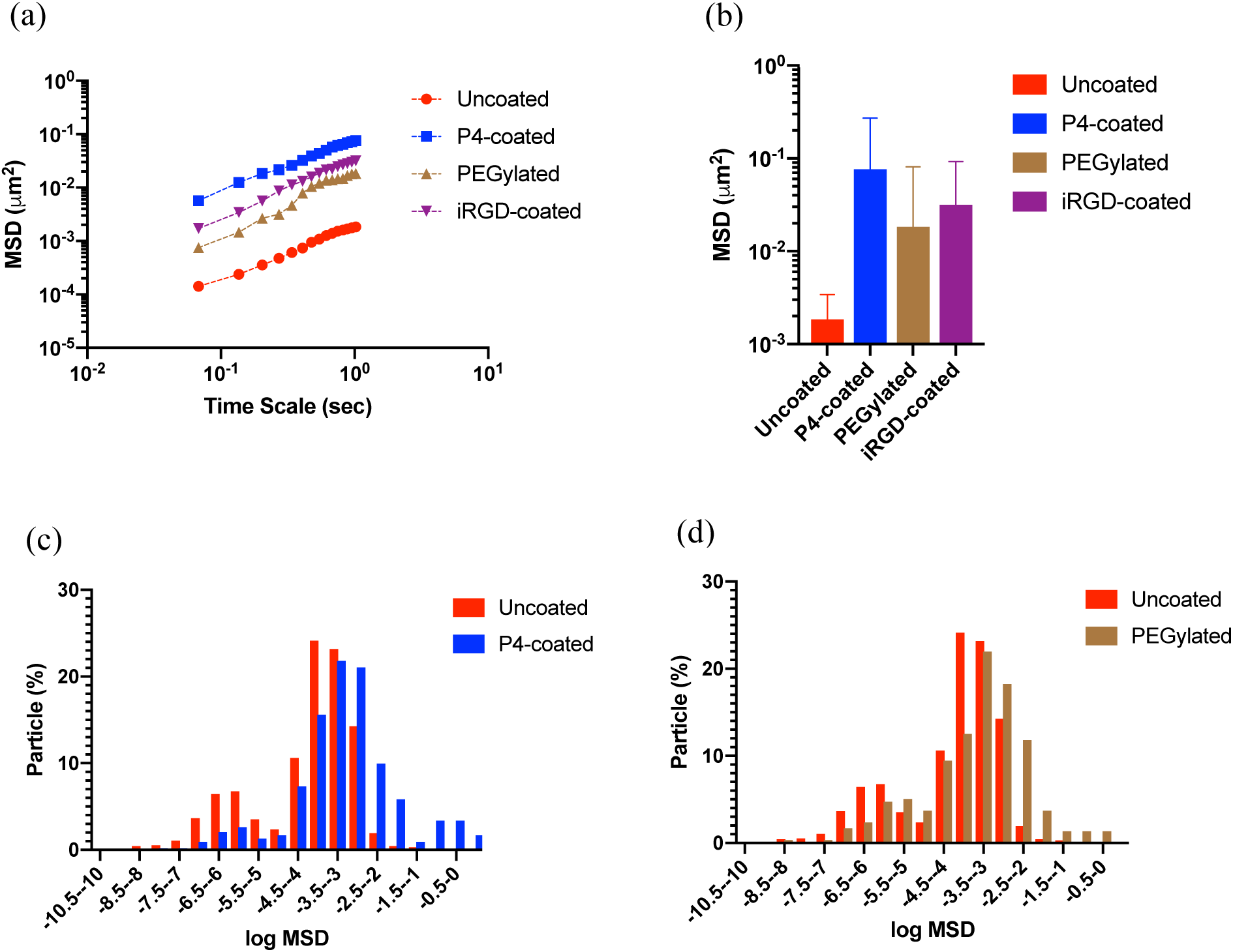
Diffusion of different surface functionalized nanoparticles through *ex vivo* tumor tissue. Ensemble-averaged mean square displacement (MSD) of peptide or PEG coated nanoparticles were compared (a) as a function of time scale and (b) at a time scale of 1 sec with the uncoated polystyrene nanoparticles using multiple particle tracking. The distribution of the logarithms of individual MSDs at a time scale of 1s of (c) uncoated versus P4-coated, and (d) uncoated vs PEGylated nanoparticles. Data represents mean ± SD for three experiments, with *n* ≥ 100 nanoparticles for each experiment.

Next, we quantified the MSD of the uncoated particle through PBS using particle tracking (**Script S1** and **Figure S4**) and used these measurements to understand how the tumor ECM hinders nanoparticle transport (**Table 3**). The ensemble-averaged MSD of the particles in tumor ECM was compared to the unhindered MSD of the uncoated bead through PBS after 1 sec time lag (**Table 3**). There was a ∼2000-fold decrease in the MSD of the uncoated nanoparticle through the tumor, whereas peptide and PEG-coatings reduced the hindrance of nanoparticle transport through the tumor tissue. P4 surface-coating most effectively reduced hindered transport, with only a ∼57-fold decrease in transport in tumor relative to unhindered transport in PBS.

## Conclusion

From combinatorial screening of a peptide-presenting phage “nanoparticle” library, we identified peptides as surface chemistries that facilitate diffusive transport through the ECM in tumor microenvironments, which can be used to ultimately improve transport of nanomedicines. Taking advantage of the molecular diversity of peptide-presenting phage libraries and their physicochemical properties, we are able to screen, iterate, and select peptides that can overcome the selection barrier of the tumor ECM to achieve permeating transport. The use of next-generation DNA sequencing allows us to expand the sequence sample space of peptides that can be analyzed and tested for improved transport. From next-generation sequencing data and *in silico* analysis, we identified specific peptides and motif sequences that enhanced transport through the ECM barrier. Select peptide-presenting clones exhibit rapid transport, as evidenced by an significant increase in their tumor transport compared to the negative control. Through systematic mutagenesis of the model P4 peptide-presenting phage, we demonstrate that alterations of the physicochemical properties and the configuration of the peptide impacts transport. We hypothesize that the mutations change the binding affinity of the peptide with the ECM, thereby hindering their diffusion. The role of binding interactions and its impact on diffusion has been observed in peptide transport in mucus and cartilage; however, further studies are warranted to confirm this mechanism in the tumor microenvironment. The model peptide-presenting phage exhibited increased uptake in tumor tissue compared to the negative control. Importantly, peptides are promising as surface coatings to improve nanoparticle transport through the tumor microenvironment. Peptide coatings reduce the hindered transport of uncoated nanoparticles through the tumor ECM and demonstrate improved transport compared to gold standard polymeric coatings. Importantly, there is a greater fraction of nanoparticles that achieve rapid transport in tumor tissues with our peptide coating compared with PEG and other controls. While additional studies are needed to study how molecular weight and density of peptide and PEG coatings affect diffusion through tumors and tumor-like environments, these findings highlight the potential of peptides as alternative coatings to improve diffusion and distribution of nanomedicines. Collectively, these results suggest that peptide-based libraries are an attractive technology that possess excellent molecular diversity from which new chemistries identified with desired functionalities to improve molecule transport and delivery through the barriers of diseases.

## Experimental

### Preparation of *in vitro* tumor ECM model

An *in vitro* tumor ECM was prepared following a method from Chauhan et al.^85^. Rat tail collagen type I at an initial concentration 8-10 mg/ml was purchased from Corning, and hyaluronic acid (HA) sodium salt isolated from rooster comb was purchased from Sigma. All subsequent steps were performed at 4 °C. HA solution was prepared by dissolving HA in 1× PBS at 5-7 mg/ml following Ramanujan et al.^67^. The solution was stirred at 350 rpm for 10 hours at 4 °C and left overnight at 4 °C. The pH and ionic strength of collagen solution were adjusted by adding 1 N NaOH and 10× PBS as recommended by the manufacturer. The HA solution was spiked into the collagen solution while stirring at 125 rpm for 10 minutes. The final concentration of HA and collagen was maintained at 0.84 mg/ml and 7.8 mg/ml, respectively, to mimic tumor ECM as reported by others^65, 67, 86^. The ECM was incubated at 37 °C for 1 hour for gelation and subsequently used for transwell or microchannel diffusion assays.

### Peptide-presenting T7 phage library screening through the *in vitro* tumor ECM

A peptide-presenting T7 phage library, which was previously developed in our lab, was used to screen and select peptide-presenting phage that permeate through the ECM. The library is a collection of phage presenting cysteine-constrained random 7-mer peptides (denoted as CX7C) on the surface of the T7 gp10 capsid. We screened the phage library against the *in vitro* tumor ECM using a transwell assay. Tumor-like ECM (thickness ∼1 mm) was prepared in the insert of a 3.0 µm pore size polyester membrane 24-well transwell (Corning). Approximately 3.2×10^8^ plaque forming units (i.e. amount of phage) of the phage library was added at the top of the tumor ECM in the donor chamber. To allow sufficient time for phage to enter ECM prior to measuring diffusivity, a time lag of 15 minutes was allotted based on theoretical calculations for phage to achieve unhindered diffusion through saline. After, the insert was transferred to a new receiving chamber filled with 600 µL of 1× PBS to maintain the hydrostatic pressure needed for diffusivity. We then performed the diffusion assay at 37 °C and collected the entire eluate from the receiving chamber after 1 hour. Part of the output was quantified, or titered, by standard double-layer plaque assay. A fraction of the eluate was used for next-generation sequencing (discussed in the next section), and the remaining fraction was amplified to make more copies that would be used for subsequent rounds of screening. The phage were amplified in BL21 *E. Coli.* (Novagen) for 3 hours as followed by the manufacturer’s recommendation and were quantified by titering. The selection was repeated for another three rounds using the same amount of input phage. With each iterative round of selection, the ECM acts a selection pressure to filter out phage clones that are not fit and cannot permeate; only clones that achieve penetrating transport through the barrier are enriched. Phage library screening were performed in replicate.

### Identification and selection of peptides using next-generation DNA sequencing

#### Sample preparation for next-generation sequencing

From each round of screening from each replicate, a fraction of the phage eluate was prepared for next-generation DNA sequencing (NGS) to identify DNA sequences that encode for the peptide-presenting phage that permeate through the ECM. Prior to NGS preparation, ten microliters of each library sample (∼10^8^ pfu/µL) was denatured at 100 °C for 15 minutes to lyse the phage and release the encapsulated DNA. The workflow for NGS sample preparation was based on manufacturers recommendation (Illumina 16S Metagenomic Sequencing Protocol). Briefly, the random insert region of the T7 library DNA was amplified from the denatured phage sample by PCR using 2× KAPA HiFi HotStart ReadyMix (KAPA Biosystems). Forward and reverse primers used for the PCR reaction were: 5’-TCG TCG GCA GCG TCA GAT GTG TAT AAG AGA CAG GTC GTA TTC CAG TCA GGT GTG-3’, and 5’-GTC TCG TGG GCT CGG AGA TGT GTA TAA GAG ACA GGA GCG CAT ATA GTT CCT CCT TTC-3’, respectively. An additional PCR was performed to attach dual indices and overhanging adapters that would be used to recognize individual samples from the final pooled library of all samples (i.e. each round from each replicate pooled together). The primers for the barcoding were provided from the Nextra XT Index kit (FC-131-1001, Illumina). The PCR amplicon was purified from the free primers using AMPure XP beads (Beckman Coulter) after each PCR amplification step. The size of the purified PCR product was verified by 2% agarose gel electrophoresis, and the DNA concentration was quantified using a Nanodrop spectrophotometer (Thermo Fisher Scientific) after each PCR step. After purification, each sample was diluted to 40 nM with 10 mM Tris buffer, pH 8.5. One microliter of each sample was pooled together, and the library was run on Illumina MiSeq (The University of Texas at Austin (UT-Austin) Genomic Sequencing and Analysis Facility).

#### Data analysis of next-generation sequencing to identify peptides

The MiSeq sequencing data was analyzed using Stampede supercomputer at the Texas Advanced Computing Center in collaboration with the Center for Computational Biology and Bioinformatics at UT-Austin. The fastq file was first demultiplexed based on the cluster’s index sequence using MiSeq Reporter, and the quality of the fastq files was checked with FastQC version 0.11.5. The fastq files were converted to fasta format using fastx-toolkit to filter out the low-quality sequence reads. The variable region encoding for the CX7C sequence was isolated using a customized Bash script. Another customized Python script was used to translate the coding DNA sequences of the randomized region to the phage-displayed peptide sequences. Here, sequences without insert, with stop codons, and/or truncations were excluded from further analysis. The total number of peptide sequences in each sample was also determined from the Python script, and the peptide sequences were sorted based on their frequency. The thirty most abundant sequences from all the samples were used for further experiments and analyses.

#### In silico physicochemical properties of the thirty most frequent peptides

Physicochemical properties such as net charge and hydropathy (i.e. hydrophobicity) of the top thirty sequences in each sample were calculated. The net charge at neutral pH of the sequences was obtained from the peptide property calculator (https://pepcalc.com/; Innovagen AB). The hydropathy, or conversely, the hydrophilicity of the peptide sequences was calculated from the grand average of hydropathy (GRAVY) score (http://www.gravy-calculator.de/)^87^. Motif analysis of the top thirty sequences in the fourth round of library screening from both replicates was done using multiple sequence alignment and visualized by Seq2logo^75^. Based on the frequency, reproducibility, and motif analysis, five sequences were selected for further validation.

### Cloning of selected peptides on T7 bacteriophage

Five peptide sequences were selected from the NGS analysis and sequence alignment to be validated for transport through tumor ECM. In addition, a negative control clone was chosen from the original library, which was not enriched with increased rounds of selection; instead, its frequency decreased with successive rounds. The peptides were each genetically engineered to be displayed on T7 based on the manufacturer’s protocol (MilliporeSigma). Briefly, the reverse complement oligonucleotides encoding each peptide sequences and the biotinylated forward oligo (Biotin-5’-ATG CTC GGG GAT CCG AAT TCT TGC-3’) were synthesized by IDT. All the oligos were dissolved in DNase free, RNase free water to a stock concentration of 100 µM and diluted to 50 µM before use. The forward and reverse oligos were annealed at 95 °C in a BioRad C1000 thermocycler (BioRad) for 3 minutes and cooled down to 25 °C at a rate of 0.1 °C/sec. The forward strand was extended from 5’ to 3’ using Klenow fragment DNA polymerase (NEB), and the extension reaction was confirmed using 10% TBE polyacrylamide gel electrophoresis. The double stranded oligonucleotide insert was double digested with HindIII and EcoRI high fidelity restriction enzymes (NEB) and ligated to the T7Select415-1b vector with precut HindIII/EcoRI overhangs (MilliporeSigma). The ligation product was packaged with the T7 packaging extracts provided by the T7Select kit (MilliporeSigma). The packaged clones were incubated with BL21 *E. coli* for infection, overlaid with liquid agar, and plated on a solid agar plate to produce plaques. Following standard techniques, individual plaques were isolated, grown in M9 liquid culture, and their DNA was purified and sequenced via Sanger DNA sequencing (UT GSAF) to validate correct insertion of the DNA sequence in the phage vector. After, the peptide-presenting phage were then amplified in liquid culture with BL21 *E. Coli.* to obtain sufficient quantities of phage needed for subsequent studies.

### Diffusion of clones through *in vitro* tumor ECM using a µ-slide assay

#### SYBR Gold labeling of phage DNA for fluorescent imaging

We wanted to fluorescently label the phage to image and determine the diffusivity through the tumor ECM. However, it was important to label the phage without affecting the coat protein, thus conserving the surface physicochemical properties of the phage clones. As a result, the encapsidated DNA of the selected peptide-presenting T7 phage was labeled with SYBR Gold dye with a slight modification of the protocol described by Mosier-Boss et al. (2003)^88^. Briefly, the SYBR Gold dye (ThermoFisher Scientific) was diluted to 100× from the 10,000× stock concentration using 1× PBS. Sixty microliters of each clone (10^8^ pfu/µL) was incubated with 500 µL of diluted dye for 20 minutes on a rocker (ThermoFisher Scientific) at room temperature. One hundred microliters of sterile 50% PEG 8000 was added to the labeled clone, incubated overnight at 4 °C, and centrifuged at 17000g for 30 minutes. The supernatant containing unlabeled free dye was decanted away from the labeled phage pellet, which was subsequently resuspended with 1× PBS. The suspension was filtered using a syringe filter 0.2 µm PES membrane (VWR). To ensure complete removal of any trace amounts of free dye, the phage suspension underwent a second round of purification using ultrafiltration with Amicon Ultra 3 kDa molecular weight cutoff of 0.5mL centrifugal filters (MilliporeSigma) and five 1× PBS washes. After centrifugation, the resulting phage pellet was resuspended with 150 µL of 1× PBS and transferred to a fresh tube.

#### Time-lapse fluorescent image acquisition in µ-slide

We visualized the transport of the fluorescently labeled phage clones along the length of a microfluidic chip over time to calculate the diffusivity of each clone, adapting from a method by Raeesi and Chan^66^. The microchannels of µ-slide (Ibidi) were loaded with 30 µL of the *in vitro* tumor ECM and incubated at 37°C for 1 hour to allow sufficient time for gelation. The channel length is 17 mm, with 3.8 mm width and 0.4 mm height. Therefore, it acts as a one-dimensional semi-infinite diffusion barrier for the labeled clones. Fifty microliters of the labeled clone were loaded into the inlet, and an equal volume of 1× PBS was loaded in the outlet reservoir to maintain the hydrostatic pressure. An Olympus IX-83 inverted fluorescence microscope with 4×/0.16NA objective, 1.63 micron/pixel image resolution and Hamamatsu ORCA-flash 4.0 camera, was used to track the fluorescent signal from the clones through the ECM. Images showing the migration of clones were acquired along the entire length and width of the channel for 15 hours with a time lapse of 30 minutes, while keeping the z-focus constant.

#### Fluorescent image analysis to determine the diffusion coefficient

The diffusion coefficient of the clones was calculated by analyzing the sequential fluorescent images using a customized Matlab script based on the procedure described elsewhere^66^. Briefly, the spatial fluorescence intensity was recorded from the average intensity over each 5-pixel penetration length of the channel (up to 1.1 mm from the input reservoir) and the entire width of the channel for all the time-lapse images. The background intensity for each channel was subtracted from the spatiotemporal intensity profile and was then normalized based on the maximum fluorescence intensity. The intensity decay was plotted along with the penetration length after 6 hours of diffusion. The average diffusion coefficient of the clones was calculated from the non-linear curve fitting of each experimental normalized intensity decay values with the one-dimensional analytical Fick’s second law of diffusion (1). The diffusion coefficient was assumed to be independent of the concentration of the clone and the penetration depth.

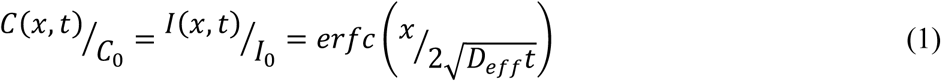

Where,

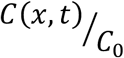: Normalized concentration profile of clone as a function of distance and time

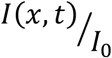: Spatio-temporal intensity profile normalized based on maximum intensity, *I*_0_

*x*: Penetration distance from the inlet along the mid-line of the channel in meters

*t*: Time of diffusion in seconds

*D*_*eff*_: Effective diffusivity of the labeled clone through tumor ECM

### Alanine mutagenesis to investigate the effect of surface physicochemical properties

Alanine mutagenesis was done on model P4 clone to understand how specific surface physicochemical properties impact diffusivity of peptide-presenting phage through tumor ECM. Specific amino acids of the peptide sequence were mutated to alanine to alter the net charge and hydropathy. We also investigated how altering the spatial distribution of physicochemical properties impacts transport by changing the amino acid position of the selected peptide sequence. Complementary oligonucleotides encoding the mutant phages were designed and synthesized by IDT. The oligonucleotides were annealed at 95 °C for 3 minutes and were cooled down to 25°C at 0.1 °C/sec using BioRad C1000 thermocycler. The annealed oligonucleotides were phosphorylated using T4 ligase buffer (NEB), and the T7Select415-1b vector arm (MilliporeSigma) was dephosphorylated following the manufacturer’s protocols. The phosphyorylated insert was ligated into the dephosphorylated vector at 16°C for 16 hours. The ligated product was then packaged, grown, and plated as described earlier. To confirm that the sequences were correctly cloned into the T7 vector, individual plaques were grown, their DNA was purified and confirmed by Sanger DNA sequencing (UT GSAF). The diffusivity of the mutants was performed and calculated using previously described microfluidic channel diffusion assays.

### Tumor tissue harvest and slice preparation for *ex vivo* study

All animal protocols were approved by the UT-Austin Institutional Animal Care and Use Committee (IACUC). Six to eight weeks old female athymic nude mice were purchased from Taconic Biosciences. Approximately 4×10^6^ cells BxPC3 cells (ATCC) in 100 µL of RPMI were mixed with 100 µL of Matrigel and subcutaneously inoculated in both flanks of mice under 2-3% isoflurane anesthesia. Tumors were palpable after three to four weeks, and mice were humanely euthanized to harvest the tumors. Tumors were removed, and tumor slices of 3 mm diameter were prepared using a reusable biopsy punch (World Precision Instruments).

### *Ex vivo* clone uptake study

The uptake of the hydrophilic, net neutral C5 clone was compared to the negative control clone following the protocol as described by Bajpayee et al.^80^. Briefly, the tumor slices were pre-equilibrated with 1× PBS supplemented with protease inhibitors (Complete ULTRA Tablets, Roche Applied Science) for 24 – 48 hours in a 37°C, 5% CO_2_ incubator. Tumor slices were then incubated with 300 µL of 10^6^ pfu/µL P4 or control-phage supplemented with protease inhibitors in a 96-well plate. The 96-well plate was incubated at 37°C, 5% CO_2_ for five days. Ten microliter samples from the bath were collected every day, and the bath titer of the clones were calculated using standard double-layer plaque assay. The number of phage were in the tumor slice was calculated based on the difference between the initial and final bath titer. At the end of 5 days, the tumor slice surfaces were blotted with Kimwipes, and wet weight of the tissue was measured. The slices were then lyophilized for 24 hours, and dry weight was measured; the water weight inside the tissue was calculated by subtracting the dry tissue weight from the wet tissue weight. The clone uptake ratio was calculated based on the clone titer inside the tissue to that of the clone titer in the bath at that given time point.

### Diffusion of surface functionalized nanoparticles in pancreatic tumor tissues

#### Surface functionalization of nanoparticles

One hundred nanometer diameter yellow-green carboxylate polystyrene nanoparticles (ThermoFisher Scientific) were surface modified with either peptide or poly(ethylene) glycol (PEG) coating. C-terminal amidated P4 and iRGD peptides (LifeTein) and methoxy-PEG 1k-amine (Alfa Aesar) were covalently conjugated to the COOH functional group of the nanoparticle at a molar ratio of 5:1 using standard carbodiimide chemistry with (1-ethyl-3-[3-dimethylaminopropyl]carbodiimide hydrochloride) (EDC) (Thermo Fisher Scientific) and Sulfo-(N-hydroxysulfosuccinimide) (Sulfo-NHS) (Thermo Fisher Scientific), following manufacturer’s protocol. After conjugation, we performed dialysis for 24 hours against 1× PBS using 100 kDa molecular weight cut-off dialysis membrane (Spectrum Labs) for buffer exchange and to remove unconjugated PEG/peptide. Size and the zeta potential of the uncoated and peptide/PEG conjugated nanoparticles were measured by suspending the particles in 10 mM NaCl at 25 °C. The nanoparticle formulations were stored at 4 °C for particle tracking experiments.

#### Nanoparticle diffusion in pancreatic tumor slices using multiple particle tracking

Diffusion of the surface functionalized nanoparticles in BxPC3 tumor tissue slices was measured by multiple particle tracking following method described in Dancy et al.^39^. Briefly, the tumor tissue slices were placed on custom microscope slide chambers, and 0.5 µL of 50 µg/mL of yellow-green fluorescent nanoparticles were injected to the *ex vivo* tumor tissue using a Hamilton syringe. The slide chambers were sealed using a coverslip, and the particles were allowed to equilibrate with the tumor tissue at 37°C for 15 minutes. The fluorescence intensity of the individual particles was then tracked at a frame rate of 15 frames/sec for 20 sec using an inverted fluorescence IX-83 Olympus microscope with 100X/1.3 NA oil-immersion objective with 65 nm/pixel image resolution. Five movies were recorded for each tissue slice, and three independent experiments were performed for each nanoparticle formulation.

Time-lapse images were analyzed using a customized Matlab script (**Script S1**) developed in our lab based on the Crocker and Grier algorithm and the Matlab version of particle tracking code repository developed by Blair & Dufresne^89, 90^. Based on the size and the threshold intensity value of the fluorescence, the centroid of the nanoparticles were determined in each image. The particle trajectory was estimated by linking the particle locations between frames, and based on a simple random-walk trajectory, the mean square displacement of the particles within a time scale was estimated. The mean square displacement (MSD) was calculated as ⟨Δ*r*^*2*^(*τ*)⟩ *=*⟨|***r***(*t* + *τ*) *-* ***r***(*t*)|^*2*^⟩, where ***r*** is the position vector of the particle, and *τ* is the lag time. The ensemble average of MSDs (⟨⟨Δ*r*^*2*^(*τ*)⟩⟩) of collected individual particles was calculated to average the MSD’s over a time lag of 1 sec over multiple particles^91^.

## Author Contributions

R.P.M. and D.G. designed experiments. R.P.M. performed experiments. X.L. assisted with the tumor injection. J.Y.K. designed the microchannel diffusion experiment and the MATLAB script, which were then modified by R.P.M. S.B. helped with the MATLAB scripts for microchannel diffusivity and particle tracking. X.P., J.L., and T.D. developed initial unpublished methods used in experiments. D.A. and D.W. developed the bioinformatics pipeline for NGS data analysis. R.P.M. and D.G. analyzed the experiments and wrote the manuscript.

## Acknowledgements

Work was supported by NIH National Heart, Lung and Blood Institute (NHLBI) of the National Institutes of Health under award number R01HL138251 and startup funds from the College of Pharmacy at University of Texas at Austin. We acknowledge The University of Texas at Austin Genome Sequencing and Analysis Facility with assistance with Sanger DNA sequencing and next-generation DNA sequencing.

## References

1. Minchinton, A. I.; Tannock, I. F., Drug penetration in solid tumours. Nat Rev Cancer 2006, 6 (8), 583–92.

2. Cilliers, C.; Menezes, B.; Nessler, I.; Linderman, J.; Thurber, G. M., Improved Tumor Penetration and Single-Cell Targeting of Antibody-Drug Conjugates Increases Anticancer Efficacy and Host Survival. Cancer research 2018, 78 (3), 758–768.

3. Ttong, R. T.; Boucher, Y.; Kozin, S. V.; Winkler, F.; Hicklin, D. J.; Jain, R. K., Vascular normalization by vascular endothelial growth factor receptor 2 blockade induces a pressure gradient across the vasculature and improves drug penetration in tumors. Cancer research 2004, 64 (11), 3731–6.

4. Koch, M.; de Jong, J. S.; Glatz, J.; Symvoulidis, P.; Lamberts, L. E.; Adams, A. L.; Kranendonk, M. E.; Terwisscha van Scheltinga, A. G.; Aichler, M.; Jansen, L.; de Vries, J.; Lub-de Hooge, M. N.; Schroder, C. P.; Jorritsma-Smit, A.; Linssen, M. D.; de Boer, E.; van der Vegt, B.; Nagengast, W. B.; Elias, S. G.; Oliveira, S.; Witkamp, A. J.; Mali, W. P.; van der Wall, E.; Garcia-Allende, P. B.; van Diest, P. J.; de Vries, E. G.; Walch, A.; van Dam, G. M.; Ntziachristos, V., Threshold Analysis and Biodistribution of Fluorescently Labeled Bevacizumab in Human Breast Cancer. Cancer research 2017, 77 (3), 623–631.

5. Chauhan, V. P.; Martin, J. D.; Liu, H.; Lacorre, D. A.; Jain, S. R.; Kozin, S. V.; Stylianopoulos, T.; Mousa, A. S.; Han, X.; Adstamongkonkul, P.; Popovic, Z.; Huang, P.; Bawendi, M. G.; Boucher, Y.; Jain, R. K., Angiotensin inhibition enhances drug delivery and potentiates chemotherapy by decompressing tumour blood vessels. Nature communications 2013, 4, 2516.

6. Primeau, A. J.; Rendon, A.; Hedley, D.; Lilge, L.; Tannock, I. F., The distribution of the anticancer drug Doxorubicin in relation to blood vessels in solid tumors. Clin Cancer Res 2005, 11 (24 Pt 1), 8782–8.

7. tannock, I. F.; Lee, C. M.; Tunggal, J. K.; Cowan, D. S.; Egorin, M. J., Limited penetration of anticancer drugs through tumor tissue: a potential cause of resistance of solid tumors to chemotherapy. Clin Cancer Res 2002, 8 (3), 878–84.

8. tunggal, J. K.; Cowan, D. S.; Shaikh, H.; Tannock, I. F., Penetration of anticancer drugs through solid tissue: a factor that limits the effectiveness of chemotherapy for solid tumors. Clin Cancer Res 1999, 5 (6), 1583–6.

9. Diop-Frimpong, B.; Chauhan, V. P.; Krane, S.; Boucher, Y.; Jain, R. K., Losartan inhibits collagen I synthesis and improves the distribution and efficacy of nanotherapeutics in tumors. Proc Natl Acad Sci U S A 2011, 108 (7), 2909–14.

10. Cabral, H.; Matsumoto, Y.; Mizuno, K.; Chen, Q.; Murakami, M.; Kimura, M.; Terada, Y.; Kano, M. R.; Miyazono, K.; Uesaka, M.; Nishiyama, N.; Kataoka, K., Accumulation of sub-100 nm polymeric micelles in poorly permeable tumours depends on size. Nat Nanotechnol 2011, 6 (12), 815–23.

11. Sykes, E. A.; Dai, Q.; Sarsons, C. D.; Chen, J.; Rocheleau, J. V.; Hwang, D. M.; Zheng, G.; Cramb, D. T.; Rinker, K. D.; Chan, W. C., Tailoring nanoparticle designs to target cancer based on tumor pathophysiology. Proc Natl Acad Sci U S A 2016, 113 (9), E1142–51.

12. Lee, J.; Kim, J.; Jeong, M.; Lee, H.; Goh, U.; Kim, H.; Kim, B.; Park, J. H., Liposome-based engineering of cells to package hydrophobic compounds in membrane vesicles for tumor penetration. Nano Lett 2015, 15 (5), 2938–44.

13. Chauhan, V. P.; Stylianopoulos, T.; Boucher, Y.; Jain, R. K., Delivery of molecular and nanoscale medicine to tumors: transport barriers and strategies. Annu Rev Chem Biomol Eng 2011, 2, 281–98.

14. Stylianopoulos, T.; Jain, R. K., Design considerations for nanotherapeutics in oncology. Nanomedicine 2015, 11 (8), 1893–907.

15. Stylianopoulos, T.; Poh, M. Z.; Insin, N.; Bawendi, M. G.; Fukumura, D.; Munn, L. L.; Jain, R. K., Diffusion of particles in the extracellular matrix: the effect of repulsive electrostatic interactions. Biophysical journal 2010, 99 (5), 1342–9.

16. Blanco, E.; Shen, H.; Ferrari, M., Principles of nanoparticle design for overcoming biological barriers to drug delivery. Nat Biotechnol 2015, 33 (9), 941–51.

17. Yuan, F.; Dellian, M.; Fukumura, D.; Leunig, M.; Berk, D. A.; Torchilin, V. P.; Jain, R. K., Vascular permeability in a human tumor xenograft: molecular size dependence and cutoff size. Cancer research 1995, 55 (17), 3752–6.

18. Hobbs, S. K.; Monsky, W. L.; Yuan, F.; Roberts, W. G.; Griffith, L.; Torchilin, V. P.; Jain, R. K., Regulation of transport pathways in tumor vessels: role of tumor type and microenvironment. Proc Natl Acad Sci U S A 1998, 95 (8), 4607–12.

19. Perrault, S. D.; Walkey, C.; Jennings, T.; Fischer, H. C.; Chan, W. C., Mediating tumor targeting efficiency of nanoparticles through design. Nano Lett 2009, 9 (5), 1909–15.

20. Popovic, Z.; Liu, W.; Chauhan, V. P.; Lee, J.; Wong, C.; Greytak, A. B.; Insin, N.; Nocera, D. G.; Fukumura, D.; Jain, R. K.; Bawendi, M. G., A nanoparticle size series for in vivo fluorescence imaging. Angew Chem Int Ed Engl 2010, 49 (46), 8649–52.

21. Huang, K.; Ma, H.; Liu, J.; Huo, S.; Kumar, A.; Wei, T.; Zhang, X.; Jin, S.; Gan, Y.; Wang, P. C.; He, S.; Zhang, X.; Liang, X. J., Size-dependent localization and penetration of ultrasmall gold nanoparticles in cancer cells, multicellular spheroids, and tumors in vivo. ACS Nano 2012, 6 (5), 4483–93.

22. Li, H. J.; Du, J. Z.; Du, X. J.; Xu, C. F.; Sun, C. Y.; Wang, H. X.; Cao, Z. T.; Yang, X. Z.; Zhu, Y. H.; Nie, S.; Wang, J., Stimuli-responsive clustered nanoparticles for improved tumor penetration and therapeutic efficacy. Proc Natl Acad Sci U S A 2016, 113 (15), 4164–9.

23. Chen, Q.; Wang, X.; Wang, C.; Feng, L.; Li, Y.; Liu, Z., Drug-Induced Self-Assembly of Modified Albumins as Nano-theranostics for Tumor-Targeted Combination Therapy. ACS Nano 2015, 9 (5), 5223–33.

24. Ruan, S.; Cao, X.; Cun, X.; Hu, G.; Zhou, Y.; Zhang, Y.; Lu, L.; He, Q.; Gao, H., Matrix metalloproteinase-sensitive size-shrinkable nanoparticles for deep tumor penetration and pH triggered doxorubicin release. Biomaterials 2015, 60, 100–10.

25. tong, R.; Chiang, H. H.; Kohane, D. S., Photoswitchable nanoparticles for in vivo cancer chemotherapy. Proc Natl Acad Sci U S A 2013, 110 (47), 19048–53.

26. tong, R.; Hemmati, H. D.; Langer, R.; Kohane, D. S., Photoswitchable nanoparticles for triggered tissue penetration and drug delivery. J Am Chem Soc 2012, 134 (21), 8848–55.

27. Wong, C.; Stylianopoulos, T.; Cui, J.; Martin, J.; Chauhan, V. P.; Jiang, W.; Popovic, Z.; Jain, R. K.; Bawendi, M. G.; Fukumura, D., Multistage nanoparticle delivery system for deep penetration into tumor tissue. Proc Natl Acad Sci U S A 2011, 108 (6), 2426–31.

28. Wang, Y.; Black, K. C.; Luehmann, H.; Li, W.; Zhang, Y.; Cai, X.; Wan, D.; Liu, S. Y.; Li, M.; Kim, P.; Li, Z. Y.; Wang, L. V.; Liu, Y.; Xia, Y., Comparison study of gold nanohexapods, nanorods, and nanocages for photothermal cancer treatment. ACS Nano 2013, 7 (3), 2068–77.

29. Chauhan, V. P.; Popovic, Z.; Chen, O.; Cui, J.; Fukumura, D.; Bawendi, M. G.; Jain, R. K., Fluorescent nanorods and nanospheres for real-time in vivo probing of nanoparticle shape-dependent tumor penetration. Angew Chem Int Ed Engl 2011, 50 (48), 11417–20.

30. Shukla, S.; DiFranco, N. A.; Wen, A. M.; Commandeur, U.; Steinmetz, N. F., To Target or Not to Target: Active vs. Passive Tumor Homing of Filamentous Nanoparticles Based on Potato virus X. Cell Mol Bioeng 2015, 8 (3), 433–444.

31. Shukla, S.; Eber, F. J.; Nagarajan, A. S.; DiFranco, N. A.; Schmidt, N.; Wen, A. M.; Eiben, S.; Twyman, R. M.; Wege, C.; Steinmetz, N. F., The Impact of Aspect Ratio on the Biodistribution and Tumor Homing of Rigid Soft-Matter Nanorods. Adv Healthc Mater 2015, 4 (6), 874–82.

32. Smith, B. R.; Kempen, P.; Bouley, D.; Xu, A.; Liu, Z.; Melosh, N.; Dai, H.; Sinclair, R.; Gambhir, S. S., Shape matters: intravital microscopy reveals surprising geometrical dependence for nanoparticles in tumor models of extravasation. Nano Lett 2012, 12 (7), 3369–77.

33. Lieleg, O.; Baumgartel, R. M.; Bausch, A. R., Selective filtering of particles by the extracellular matrix: an electrostatic bandpass. Biophysical journal 2009, 97 (6), 1569–77.

34. Wang, H. X., Zuo, Z.Q., Du, J.Z., Wang, Y.C., Sun, R., Cao, Z.T., Ye, X.D., Wang, J.L., Leong, K.W., Wang, J., Surface charge critically affects tumor penetration and therapeutic efficacy of cancer nanomedicines. Nano Today 2016, 11 (2), 133–144.

35. Wang, G.; Chen, Y.; Wang, P.; Wang, Y.; Hong, H.; Li, Y.; Qian, J.; Yuan, Y.; Yu, B.; Liu, C., Preferential tumor accumulation and desirable interstitial penetration of poly(lacticco-glycolic acid) nanoparticles with dual coating of chitosan oligosaccharide and polyethylene glycol-poly(D,L-lactic acid). Acta Biomater 2016, 29, 248–260.

36. Campbell, R. B.; Fukumura, D.; Brown, E. B.; Mazzola, L. M.; Izumi, Y.; Jain, R. K.; Torchilin, V. P.; Munn, L. L., Cationic charge determines the distribution of liposomes between the vascular and extravascular compartments of tumors. Cancer research 2002, 62 (23), 6831–6.

37. Jin, S.; Ma, X.; Ma, H.; Zheng, K.; Liu, J.; Hou, S.; Meng, J.; Wang, P. C.; Wu, X.; Liang, X. J., Surface chemistry-mediated penetration and gold nanorod thermotherapy in multicellular tumor spheroids. Nanoscale 2013, 5 (1), 143–6.

38. tomasetti, L.; Liebl, R.; Wastl, D. S.; Breunig, M., Influence of PEGylation on nanoparticle mobility in different models of the extracellular matrix. European journal of pharmaceutics and biopharmaceutics: official journal of Arbeitsgemeinschaft fur Pharmazeutische Verfahrenstechnik e.V 2016, 108, 145–155.

39. Dancy, J. G.; Wadajkar, A. S.; Schneider, C. S.; Mauban, J. R. H.; Goloubeva, O. G.; Woodworth, G. F.; Winkles, J. A.; Kim, A. J., Non-specific binding and steric hindrance thresholds for penetration of particulate drug carriers within tumor tissue. Journal of controlled release: official journal of the Controlled Release Society 2016, 238, 139–148.

40. Nance, E.; Zhang, C.; Shih, T. Y.; Xu, Q.; Schuster, B. S.; Hanes, J., Brain-penetrating nanoparticles improve paclitaxel efficacy in malignant glioma following local administration. ACS Nano 2014, 8 (10), 10655–64.

41. Nance, E. A.; Woodworth, G. F.; Sailor, K. A.; Shih, T. Y.; Xu, Q.; Swaminathan, G.; Xiang, D.; Eberhart, C.; Hanes, J., A dense poly(ethylene glycol) coating improves penetration of large polymeric nanoparticles within brain tissue. Science translational medicine 2012, 4 (149), 149ra119.

42. Nance, E.; Timbie, K.; Miller, G. W.; Song, J.; Louttit, C.; Klibanov, A. L.; Shih, T. Y.; Swaminathan, G.; Tamargo, R. J.; Woodworth, G. F.; Hanes, J.; Price, R. J., Non-invasive delivery of stealth, brain-penetrating nanoparticles across the blood-brain barrier using MRI-guided focused ultrasound. Journal of controlled release: official journal of the Controlled Release Society 2014, 189, 123–132.

43. Mishra, S.; Webster, P.; Davis, M. E., PEGylation significantly affects cellular uptake and intracellular trafficking of non-viral gene delivery particles. Eur J Cell Biol 2004, 83 (3), 97–111.

44. Klibanov, A. L.; Maruyama, K.; Beckerleg, A. M.; Torchilin, V. P.; Huang, L., Activity of amphipathic poly(ethylene glycol) 5000 to prolong the circulation time of liposomes depends on the liposome size and is unfavorable for immunoliposome binding to target. Biochim Biophys Acta 1991, 1062 (2), 142–8.

45. Betker, J. L.; Gomez, J.; Anchordoquy, T. J., The effects of lipoplex formulation variables on the protein corona and comparisons with in vitro transfection efficiency. Journal of controlled release: official journal of the Controlled Release Society 2013, 171 (3), 261–8.

46. Remaut, K.; Lucas, B.; Braeckmans, K.; Demeester, J.; de Smedt, S. C., Pegylation of liposomes favours the endosomal degradation of the delivered phosphodiester oligonucleotides. Journal of controlled release: official journal of the Controlled Release Society 2007, 117 (2), 256–66.

47. Ganson, N. J.; Povsic, T. J.; Sullenger, B. A.; Alexander, J. H.; Zelenkofske, S. L.; Sailstad, J. M.; Rusconi, C. P.; Hershfield, M. S., Pre-existing anti-polyethylene glycol antibody linked to first-exposure allergic reactions to pegnivacogin, a PEGylated RNA aptamer. J Allergy Clin Immunol 2016, 137 (5), 1610–1613 e7.

48. Garay, R. P.; El-Gewely, R.; Armstrong, J. K.; Garratty, G.; Richette, P., Antibodies against polyethylene glycol in healthy subjects and in patients treated with PEG-conjugated agents. Expert Opin Drug Deliv 2012, 9 (11), 1319–23.

49. Hershfield, M. S.; Ganson, N. J.; Kelly, S. J.; Scarlett, E. L.; Jaggers, D. A.; Sundy, J. S., Induced and pre-existing anti-polyethylene glycol antibody in a trial of every 3-week dosing of pegloticase for refractory gout, including in organ transplant recipients. Arthritis Res Ther 2014, 16 (2), R63.

50. Armstrong, J. K.; Hempel, G.; Koling, S.; Chan, L. S.; Fisher, T.; Meiselman, H. J.; Garratty, G., Antibody against poly(ethylene glycol) adversely affects PEG-asparaginase therapy in acute lymphoblastic leukemia patients. Cancer 2007, 110 (1), 103–11.

51. Verhoef, J. J.; Carpenter, J. F.; Anchordoquy, T. J.; Schellekens, H., Potential induction of anti-PEG antibodies and complement activation toward PEGylated therapeutics. Drug Discov Today 2014, 19 (12), 1945–52.

52. Mima, Y.; Hashimoto, Y.; Shimizu, T.; Kiwada, H.; Ishida, T., Anti-PEG IgM Is a Major Contributor to the Accelerated Blood Clearance of Polyethylene Glycol-Conjugated Protein. Molecular pharmaceutics 2015, 12 (7), 2429–35.

53. Ishida, T.; Kiwada, H., Accelerated blood clearance (ABC) phenomenon upon repeated injection of PEGylated liposomes. International journal of pharmaceutics 2008, 354 (1-2), 56–62.

54. Keefe, A. J.; Caldwell, K. B.; Nowinski, A. K.; White, A. D.; Thakkar, A.; Jiang, S., Screening nonspecific interactions of peptides without background interference. Biomaterials 2013, 34 (8), 1871–7.

55. Ghosh, D.; Peng, X.; Leal, J.; Mohanty, R., Peptides as drug delivery vehicles across biological barriers. J Pharm Investig 2018, 48 (1), 89–111.

56. Komin, A.; Russell, L. M.; Hristova, K. A.; Searson, P. C., Peptide-based strategies for enhanced cell uptake, transcellular transport, and circulation: Mechanisms and challenges. Adv Drug Deliv Rev 2017, 110-111, 52–64.

57. Ruoslahti, E., Tumor penetrating peptides for improved drug delivery. Adv Drug Deliv Rev 2017, 110-111, 3–12.

58. Pasqualini, R., Vascular targeting with phage peptide libraries. The quarterly journal of nuclear medicine: official publication of the Italian Association of Nuclear Medicine (AIMN) [and] the International Association of Radiopharmacology (IAR) 1999, 43 (2), 159–62.

59. Sugahara, K. N.; Teesalu, T.; Karmali, P. P.; Kotamraju, V. R.; Agemy, L.; Girard, O. M.; Hanahan, D.; Mattrey, R. F.; Ruoslahti, E., Tissue-penetrating delivery of compounds and nanoparticles into tumors. Cancer cell 2009, 16 (6), 510–20.

60. Shao, Q.; Jiang, S., Molecular understanding and design of zwitterionic materials. Adv Mater 2015, 27 (1), 15–26.

61. Chen, S.; Cao, Z.; Jiang, S., Ultra-low fouling peptide surfaces derived from natural amino acids. Biomaterials 2009, 30 (29), 5892–6.

62. Liu, E. J.; Sinclair, A.; Keefe, A. J.; Nannenga, B. L.; Coyle, B. L.; Baneyx, F.; Jiang, S., EKylation: Addition of an Alternating-Charge Peptide Stabilizes Proteins. Biomacromolecules 2015, 16 (10), 3357–61.

63. Li, L. D.; Crouzier, T.; Sarkar, A.; Dunphy, L.; Han, J.; Ribbeck, K., Spatial configuration and composition of charge modulates transport into a mucin hydrogel barrier. Biophysical journal 2013, 105 (6), 1357–65.

64. Weiss, G. A., Editorial overview: How to generate molecular diversity, the most important process in biology. Curr Opin Chem Biol 2017, 41, A3–A5.

65. DuFort, C. C.; DelGiorno, K. E.; Carlson, M. A.; Osgood, R. J.; Zhao, C.; Huang, Z.; Thompson, C. B.; Connor, R. J.; Thanos, C. D.; Scott Brockenbrough, J.; Provenzano, P. P.; Frost, G. I.; Michael Shepard, H.; Hingorani, S. R., Interstitial Pressure in Pancreatic Ductal Adenocarcinoma Is Dominated by a Gel-Fluid Phase. Biophysical journal 2016, 110 (9), 2106–19.

66. Raeesi, V.; Chan, W. C., Improving nanoparticle diffusion through tumor collagen matrix by photo-thermal gold nanorods. Nanoscale 2016, 8 (25), 12524–30.

67. Ramanujan, S.; Pluen, A.; McKee, T. D.; Brown, E. B.; Boucher, Y.; Jain, R. K., Diffusion and convection in collagen gels: implications for transport in the tumor interstitium. Biophysical journal 2002, 83 (3), 1650–60.

68. Aoki, S.; Morohashi, K.; Sunoki, T.; Kuramochi, K.; Kobayashi, S.; Sugawara, F., Screening of paclitaxel-binding molecules from a library of random peptides displayed on T7 phage particles using paclitaxel-photoimmobilized resin. Bioconjugate chemistry 2007, 18 (6), 1981–6.

69. Liu, G. W.; Livesay, B. R.; Kacherovsky, N. A.; Cieslewicz, M.; Lutz, E.; Waalkes, A.; Jensen, M. C.; Salipante, S. J.; Pun, S. H., Efficient Identification of Murine M2 Macrophage Peptide Targeting Ligands by Phage Display and Next-Generation Sequencing. Bioconjugate chemistry 2015, 26 (8), 1811–7.

70. Dennis, M. S., Selection and Screening Strategies. In Phage Display in Biotechnology and Drug Delivery, Second Edition ed.; Sachdev S. Sidhu, C. R. G., Ed. CRC Press: Boca Raton, 2015; pp 97–111.

71. Scott, J. K.; Smith, G. P., Searching for peptide ligands with an epitope library. Science 1990, 249 (4967), 386–90.

72. Barry, M. A.; Dower, W. J.; Johnston, S. A., Toward cell-targeting gene therapy vectors: selection of cell-binding peptides from random peptide-presenting phage libraries. Nature medicine 1996, 2 (3), 299–305.

73. Matochko, W. L.; Cory Li, S.; Tang, S. K.; Derda, R., Prospective identification of parasitic sequences in phage display screens. Nucleic Acids Res 2014, 42 (3), 1784–98.

74. Ravn, U.; Didelot, G.; Venet, S.; Ng, K. T.; Gueneau, F.; Rousseau, F.; Calloud, S.; Kosco-Vilbois, M.; Fischer, N., Deep sequencing of phage display libraries to support antibody discovery. Methods (San Diego, Calif.) 2013, 60 (1), 99–110.

75. thomsen, M. C.; Nielsen, M., Seq2Logo: a method for construction and visualization of amino acid binding motifs and sequence profiles including sequence weighting, pseudo counts and two-sided representation of amino acid enrichment and depletion. Nucleic acids research 2012, 40 (Web Server issue), W281–7.

76. Sunderland, K. S.; Yang, M.; Mao, C., Phage-Enabled Nanomedicine: From Probes to Therapeutics in Precision Medicine. Angewandte Chemie (International ed. in English) 2017, 56 (8), 1964–1992.

77. Hu, J.; Miyanaga, K.; Tanji, Y., Diffusion of bacteriophages through artificial biofilm models. Biotechnology progress 2012, 28 (2), 319–26.

78. Hu, J.; Miyanaga, K.; Tanji, Y., Diffusion properties of bacteriophages through agarose gel membrane. Biotechnol Prog 2010, 26 (5), 1213–21.

79. Lohr, M.; Trautmann, B.; Gottler, M.; Peters, S.; Zauner, I.; Maillet, B.; Kloppel, G., Human ductal adenocarcinomas of the pancreas express extracellular matrix proteins. Br J Cancer 1994, 69 (1), 144–51.

80. Bajpayee, A. G.; Wong, C. R.; Bawendi, M. G.; Frank, E. H.; Grodzinsky, A. J., Avidin as a model for charge driven transport into cartilage and drug delivery for treating early stage post-traumatic osteoarthritis. Biomaterials 2014, 35 (1), 538–49.

81. Bajpayee, A. G.; De la Vega, R. E.; Scheu, M.; Varady, N. H.; Yannatos, I. A.; Brown, L. A.; Krishnan, Y.; Fitzsimons, T. J.; Bhattacharya, P.; Frank, E. H.; Grodzinsky, A. J.; Porter, R. M., Sustained intra-cartilage delivery of low dose dexamethasone using a cationic carrier for treatment of post traumatic osteoarthritis. Eur Cell Mater 2017, 34, 341–364.

82. Bajpayee, A. G.; Scheu, M.; Grodzinsky, A. J.; Porter, R. M., Electrostatic interactions enable rapid penetration, enhanced uptake and retention of intra-articular injected avidin in rat knee joints. J Orthop Res 2014, 32 (8), 1044–51.

83. Lai, S. K.; O’Hanlon, D. E.; Harrold, S.; Man, S. T.; Wang, Y. Y.; Cone, R.; Hanes, J., Rapid transport of large polymeric nanoparticles in fresh undiluted human mucus. Proceedings of the National Academy of Sciences of the United States of America 2007, 104 (5), 1482–7.

84. Xu, Q.; Ensign, L. M.; Boylan, N. J.; Schon, A.; Gong, X.; Yang, J. C.; Lamb, N. W.; Cai, S.; Yu, T.; Freire, E.; Hanes, J., Impact of Surface Polyethylene Glycol (PEG) Density on Biodegradable Nanoparticle Transport in Mucus ex Vivo and Distribution in Vivo. ACS nano 2015, 9 (9), 9217–27.

85. Chauhan, V. P.; Lanning, R. M.; Diop-Frimpong, B.; Mok, W.; Brown, E. B.; Padera, T. P.; Boucher, Y.; Jain, R. K., Multiscale measurements distinguish cellular and interstitial hindrances to diffusion in vivo. Biophysical journal 2009, 97 (1), 330–6.

86. Netti, P. A.; Berk, D. A.; Swartz, M. A.; Grodzinsky, A. J.; Jain, R. K., Role of extracellular matrix assembly in interstitial transport in solid tumors. Cancer research 2000, 60 (9), 2497–503.

87. Kyte, J.; Doolittle, R. F., A simple method for displaying the hydropathic character of a protein. Journal of molecular biology 1982, 157 (1), 105–32.

88. Mosier-Boss, P. A.; Lieberman, S. H.; Andrews, J. M.; Rohwer, F. L.; Wegley, L. E.; Breitbart, M., Use of fluorescently labeled phage in the detection and identification of bacterial species. Applied spectroscopy 2003, 57 (9), 1138–44.

89. Crocker, J. C.; Grier, D. G., Methods of Digital Video Microscopy for Colloidal Studies. Journal of Colloid and Interface Science 1996, 179 (1), 298–310.

90. Dufresne, D. B. a. E. The Matlab Particle Tracking Code Repository. http://site.physics.georgetown.edu/matlab/index.html.

91. Schuster, B. S.; Ensign, L. M.; Allan, D. B.; Suk, J. S.; Hanes, J., Particle tracking in drug and gene delivery research: State-of-the-art applications and methods. Advanced drug delivery reviews 2015, 91, 70–91.

