## Supplemental Information for "Identification of peptide coatings that enhance diffusive transport of nanoparticles through the tumor microenvironment"

Debadyuti Ghosh

(512) 232-7155

**Supplementary Note 1. In silico analysis evaluates the overall physicochemical properties of the thirty most frequent peptide sequences**

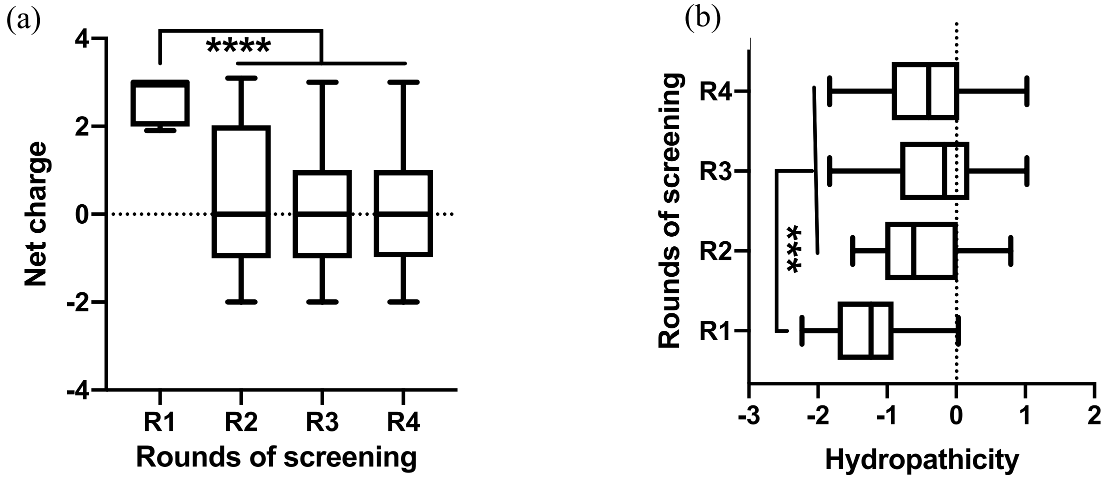


**Figure S1. Physicochemical properties of phage after selection rounds.** Average (a) net charge at pH 7 and (b) hydropathicity of the top thirty frequent sequences diffused over four rounds of screening (R1, R2, R3, and R4) analyzed from NGS data. Data represents median, IQR. One-way analysis of variance (ANOVA) with Dunnett's T3 multiple comparisons test ($p<0.0001$).

The net charge of the top thirty frequent peptides from each round of screening was calculated, and their overall net charge was plotted (**Figure S1a**). From the whisker plot, the sequences from the first round (denoted as R1) have a median charge of +3, and all analyzed sequences were basic. In subsequent rounds (denoted as R2 – R4), there is a decrease in the median net charge compared to R1 ($p < 0.0001$). The median net charge changes from +3 in R1 to 0 in R2 to R4, and this finding suggests that during screening, there is a selection for net neutral charge peptides that could permeate through the *in vitro* tumor ECM. While the tumor ECM has overall net negative charge, the collagen fibers has a slightly positive surface charge density of 0.002 $C/m^2$ and hyaluronic acid has a highly negative surface charge density of -0.10 $C/m^2$ [15]. Presence of both of these components may result in localized charge patches, which makes the ECM an effective and complex electrostatic filter that is able to restrict the diffusion of both positively and negatively charged molecules[33, 68-69]. Due to the role of these electrostatic interactions, neutral molecules can penetrate more rapidly compared to their charged counterparts.

Next, we characterized the overall hydrophobicity of our most abundant sequences. The Grand Average of Hydropathy (GRAVY), which is a score of average hydrophobicity estimated from an amino acid sequence, was calculated for each peptide from the top thirty sequences in each round and replicate (**Figure S1b**)[70]. A negative GRAVY score represents an amino acid sequence that is hydrophilic, whereas a positive score indicates the amino acid sequence is hydrophobic[71]. Most of peptides from the R1 eluate are hydrophilic, ranging from a minimum hydropathicity score of -2.23 to a maximum of 0.03. With each successive round, there is an increase in the minimum and maximum GRAVY score; the minimum and maximum hydropathy scores range from -1.83 to 1.02 in the R3 and R4 eluates. The net median hydropathy increases significantly from -1.23 in R1 to -0.4 in R4 ($p = 0.0002$). This suggests there are less hydrophilic amino acids of selected peptides that permeate through the tumor-like ECM, but generally the sequences are net hydrophilic. This change could be attributed to the loss of basic amino acids during selection and the increased presence of glycines. Glycines have been used as flexible linkers to provide greater conformational stability; however they contribute to the hydrophobicity of a sequence. Selection of net neutral charged peptides over rounds of screening (**Figure S1a**) also indicates a reduction in the number of both acidic and basic amino acid residues. Reduction in these amino acids simultaneously resulted in the decrease of the net hydrophilicity of the peptides in higher rounds.

**Table S1. Diffusivity of different T7 phage in PBS measured by dynamic light scattering**

Unhindered diffusivity of phage clones was calculated from dynamic light scattering (DLS) measurement in 1× PBS at 25 °C (Zetasizer Nano, Malvern Instruments). Clones of $10^7$-$10^8$ pfu/μL titer were suspended in 1× PBS and syringe filtered with a 0.2 μm PES membrane (VWR) to remove any remaining bacteria debris before the measurement.

| **Peptide Name** | **Diffusivity in PBS at 25°C ($m^2$/sec)** |
|---|---|
| WT | $(5.30\pm0.09)\times10^{-12}$ |
| P1-phage | $(4.82\pm0.02)\times10^{-12}$ |
| P2-phage | $(6.27\pm0.12)\times10^{-12}$ |
| P3-phage | $(6.07\pm0.01)\times10^{-12}$ |
| P4-phage | $(6.06\ \pm0.02)\times10^{-12}$ |
| P5-phage | $(5.74\ \pm0.01)\times10^{-12}$ |
| Control-phage | $(5.75\pm0.08)\times10^{-12}$ |

**Determination of diffusion coefficient of clones through tumor ECM using microchannel diffusion assay**

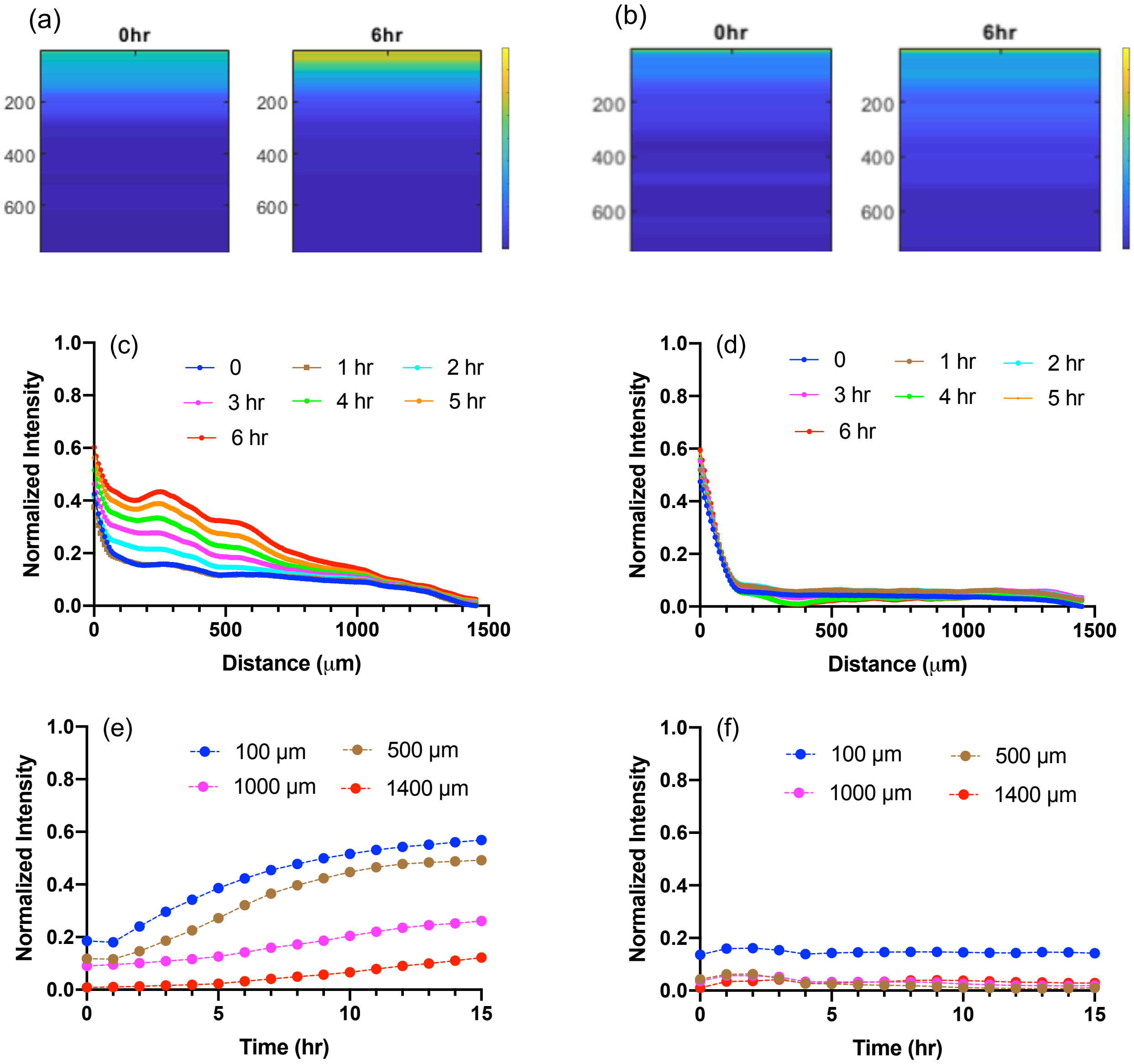


**Figure S2.** Representative images obtained from MATLAB analysis for μ-slide diffusion assay are compared for P4-phage and the negative control-phage. The colormaps generated in Matlab are showing the intensity profile of the fluorescently-labeled (a) P4-phage and (b) negative control-phage at initial time point (time 0) and after 6 hours of diffusion through the microchannel. P4-phage penetrates a longer distance in 6 hours compared to the control-phage. The normalized fluorescent intensity is plotted against the channel length from the input reservoir up to 6 hours with 1 hour time interval for (c) P4-phage and (d) control-phage. Although at the inlet reservoir the normalized intensity is ~0.6 for both of the clones initially, the normalized intensity dropped to less than 0.1 for the negative control-phage within 100 µm, and it remained the same even at higher time points. However, P4-phage penetrated up to ~1.45 mm of the channel length and the intensity increased with time for each penetration depth. Similarly, the evolution of normalized intensity with time is studied at fixed penetration distance up to initial 1.4 mm for (e) P4-phage and (f) control-phage. The normalized intensity increases slightly in the first 1 hour, but remained constant throughout the higher time points up to 15 hours, and the value remained within 0.2 throughout for the control-phage. On the other hand, the normalized intensity increased steadily to 0.6 with time for all the channel lengths reported for P4-phage.

**Supplementary Note 2. Hydrophilic and net neutral clone, C4 diffuses faster than other selected clones through tumor ECM using a transwell assay**

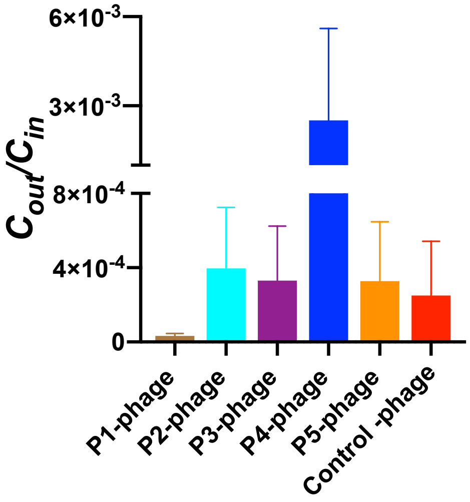

**Figure S3.** Higher output titer ($C_{out}$) of the clone, P4-phage is collected after 1 hr. of diffusion through the transwell compared to other selected clones and the negative control-phage while keeping the phage input ($C_{in}$) to be the same.

Diffusion of the selected clones is validated using bulk transwell diffusion assay in which the clones are made to permeate through the *in vitro* tumor ECM. A 1mm thick tumor ECM is prepared in the insert of a 3.0 μm pore size polyester membrane 24-well transwell (Corning) as described in the phage library screening section. 70 μL of $4.6\times10^{6}$ pfu/μL clone is added at the top of the ECM as input in the donor chamber of the transwell. The clones are rested on the ECM for 15 minutes before the diffusion assay to account for the unhindered diffusion of the clones because of the ECM leakiness. Then the insert is transferred to a new receiving chamber filled with 600 μL of PBS to equilibrate the hydrostatic pressure and allow the clone to diffuse through the ECM. 50 μL samples are collected at time intervals of 10, 20, 30, 40 and 50 minutes, and an equal volume of fresh PBS 1× is replenished. Temperature is maintained at 37 °C throughout the diffusion experiment. After 1 hour, all the eluates from the receiving chamber is collected. The output titer of the clones is quantified using standard double-layer plaque assay to compare the output titer of the clones and the negative control-phage. Comparison of the output titer per the constant input titer (**Figure S3**) shows a hydrophilic and net neutral surface charge clone, P4-phage has higher penetration through the tumor ECM compared to all the other selected clones and the control-phage.

**Script S1.** MATLAB script for Particle tracking

```
clear all
%create cells to store data for all the images in the file
```

```
I=cell(1,300);
J=cell(1,300);
b=cell(1,300);
pk=cell(1,300);

%Read images
for i=1:300
    if i>=1 && i<=9
    I{i}=double(imread(sprintf('Image_T000%d.tif',i)));
    elseif i>=10 && i<=99
            I{i}=double(imread(sprintf('Image_T00%d.tif',i)));
    elseif  i>=100 && i<=300
            I{i}=double(imread(sprintf('Image_T0%d.tif',i)));
    end
    %doing this because RGB image, last column access different colors
    J{i}(:,:)=I{i}(:,:,1);
    whos J{i};
    %colormap('gray'), imagesc(J)
    b{i}=bpass(J{i},0,size of particle,cut-off intensity);
    pk{i}=pkfnd(b{i},minimum value of intensity of local maxima, size)
    max(max(b{i}));
    % N x 4 array containing, x, y and brightness for each feature
    % out(:,1) is the x-coordinates
    % out(:,2) is the y-coordinates
    % out(:,3) is the brightnesses
    % out(:,4) is the sqare of the radius of gyration
    cnt{i}=cntrd(b{i},pk{i},size+2);
    lencnt(1,i)=size(cnt{i},1);
end
```

```
lencnt2=lencnt';


%Extract only x- and y-coordinates of the particles

B= cell2mat(cnt')

sizB=size(B,2);

k=0;

for i=1:size(lencnt2,1)

    if i~=1

    k=k+lencnt2(i-1);

    end

    B(k+1:k+lencnt2(i),sizB+1)=i;

end

B


%Tracking of particles

tr = track(B, maximum displacement)

T= table(tr);

%Export tracking information to excel

filename='trackeddata.xlsx';

writetable(T,filename,'Sheet',1)
```

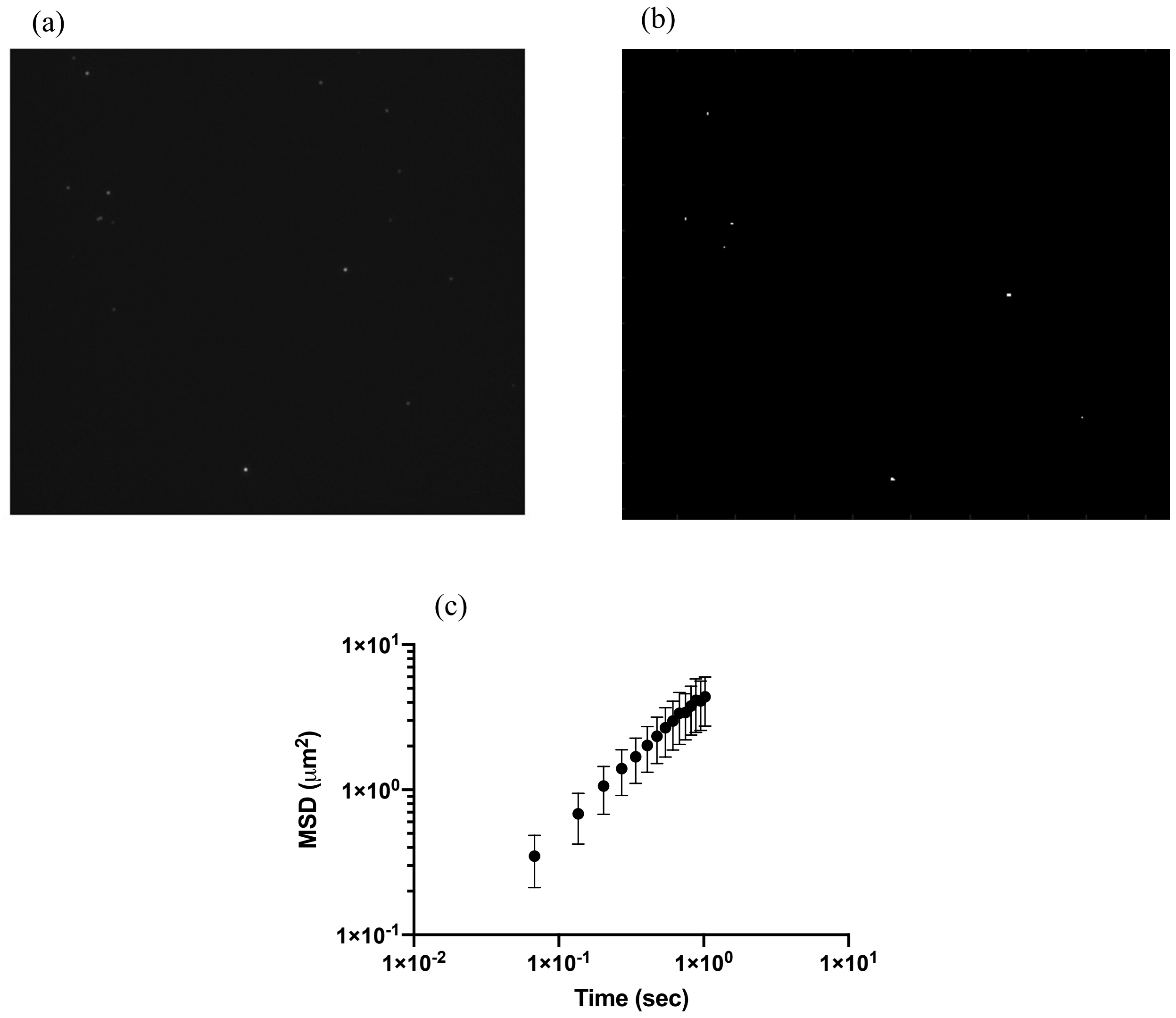


**Figure S4.** (a) Representative image (in grey-scale) obtained from the time-lapse particle tracking of uncoated polystyrene nanoparticle in PBS. (b) Matlab processed image assuming size 14 pixel (estimated from ImageJ) and cut-off intensity 100. (c) Ensemble-averaged mean square displacement (MSD) of the uncoated polystyrene nanoparticle in PBS as a function of time.
